# Metal ion binding of vimentin tail domain fragments

**DOI:** 10.1101/2025.04.10.648257

**Authors:** Estely J. Carranza, Marie C. Heffern, Dylan T. Murray

## Abstract

The intermediate filament (IF) protein vimentin is widely distributed in various cell types in the body and is vital for the proper maintenance of the cell cytoskeletal architecture, yet an extensive structural characterization of its head and tail domains remains elusive. Alterations in the assembly and organization of vimentin IFs, including filament network reorganization, have been associated with several diseases including cataracts, myopathies, and metastatic cancer. The C-terminal tail domain of vimentin is of increasing interest as it is essential for regulating the structure and mechanical properties of filament networks through interactions with divalent metal ions, but the molecular basis of these tail domain-metal interactions have not been characterized. In this work, we perform an in-depth analysis of the structural and metal binding properties of fragments of the vimentin tail domain. Mass spectrometry, and UV-Vis and circular dichroism (CD) spectroscopy reveal the direct binding of divalent copper (Cu(II)) to the last 11 residues of the tail domain. Solution nuclear magnetic resonance (NMR) and CD measurements show that in isolation, the complete vimentin tail domain is primarily disordered, and that Cu(II)-binding involves both the last 11 residues and another segment in the middle of the tail domain. Aside from these binding sites, Cu(II) does not induce any significant ordering of the tail domain. These findings further support the tail domain serving as a key metal binding region of vimentin and provide new insights into the important interplay between the tail domain and metals in vimentin IF physiology and pathology.

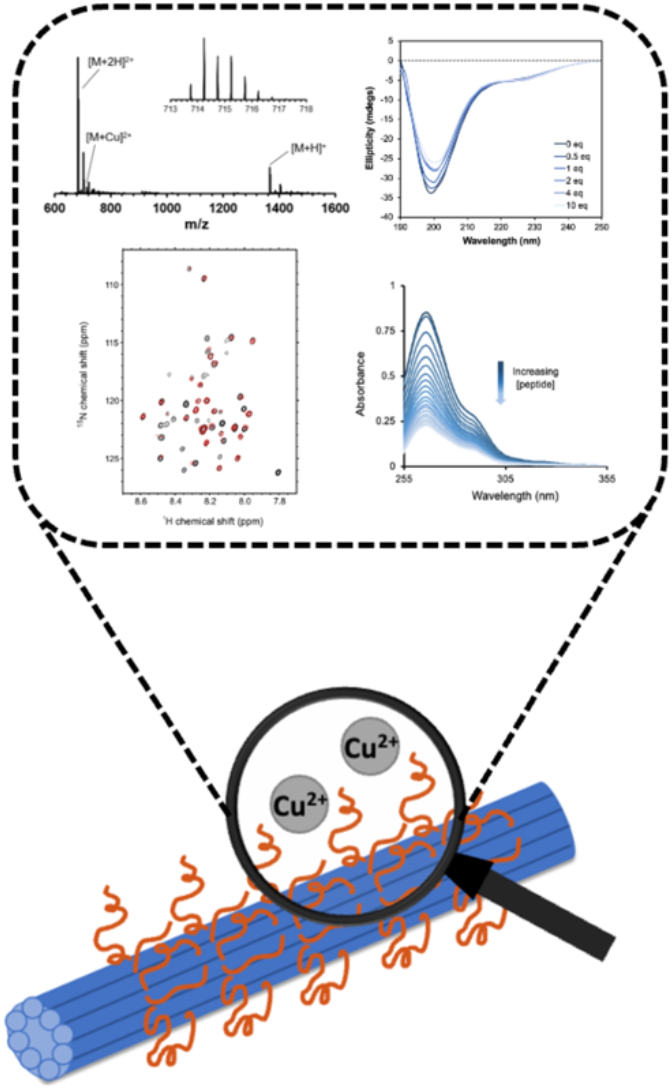

## Introduction

Vimentin intermediate filaments (IFs) are vital for the proper maintenance of cytoskeletal architecture and dynamics. Vimentin performs critical functions in cell adhesion, migration, and signaling and is considered an active agent and marker of the epithelial to mesenchymal transition, a key process in cancer metastasis.^1,2^ Alterations in the expression, structure, and organization of vimentin IFs have been linked to numerous diseases including cataracts, myopathies, and epithelial cancers.^3^ However, the underlying molecular mechanisms that drive the dysregulation and disruption of mature vimentin filament structures in disease and how these structures differ from normal physiological forms, remain uncharacterized.

Vimentin monomers have a three-part domain structure, consisting of a highly ordered central rod domain, and intrinsically disordered N-terminal head and C-terminal tail domains (Figure 1A). The macroscopic, hierarchical assembly process of vimentin has been well-described, involving the formation of dimers that undergo several self-association steps to assemble into the extended, mature filaments.^4,5^ Previous structural studies have elucidated the structural features and role of the head domain in filament formation,^6–9^ led to the complete structure determination of the rod domain,^10–15^ and have revealed insights into the molecular structure of oligomeric forms of vimentin.^5,16,17^ Recently, cryo-electron tomography and single particle methods were applied to study the cellular structures of vimentin IFs providing a 7.2 Å model of mature, cytoskeletal vimentin and demonstrated the importance of regions of low-complexity (head and tail domains) in the overall structure of IFs.^18^ However, a high-resolution atomic-level understanding of the complete architecture of vimentin IF assemblies, particularly of the dynamic properties of its disordered head and tail regions, is largely missing.^19^

**Figure 1.**
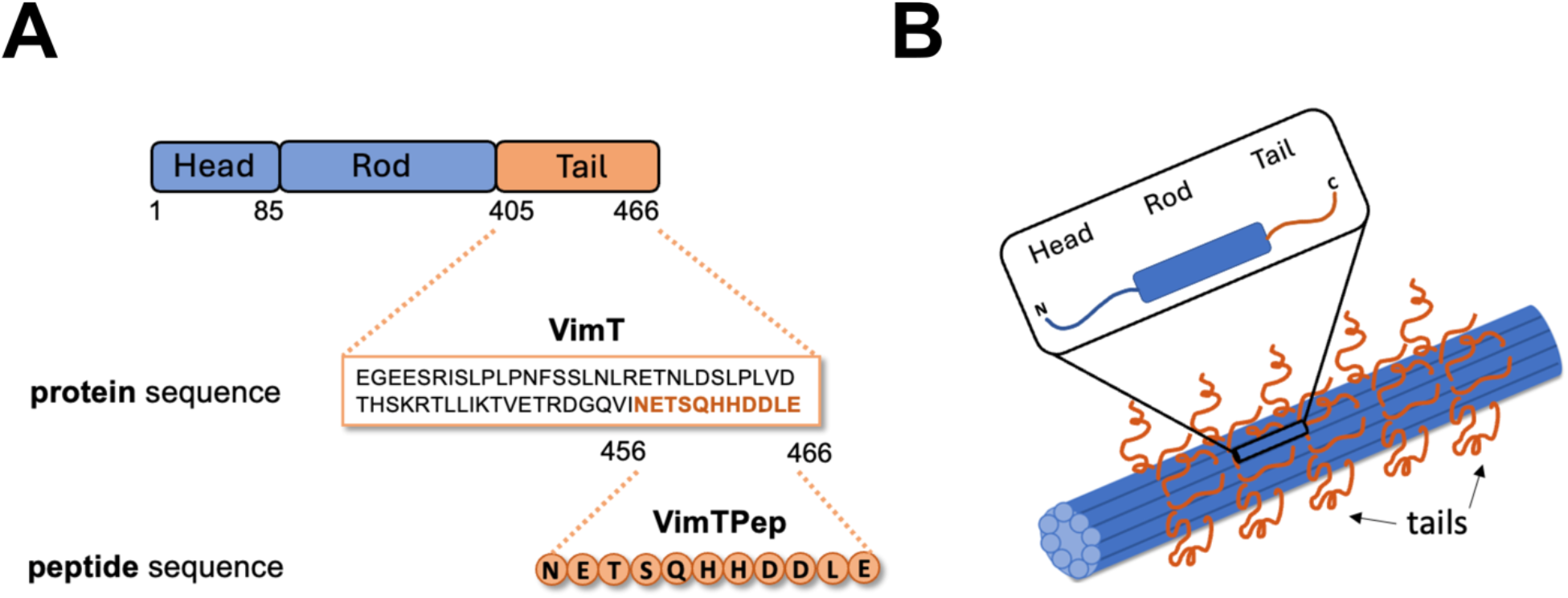
Vimentin protein domains and filament organization. (A) Diagram of all three domains of vimentin highlighting the tail domain regions investigated in this study. (B) The proposed bottlebrush-like arrangement of vimentin tails along the filament core, with a zoomed in view of the monomer units that come together to assemble into the extended filament.^34^

As a highly negatively charged biopolymer, metal ions modulate vimentin self-association and filament organization, and influences the mechanical properties of vimentin assemblies.^20–22^ *In vitro*, vimentin IF assembly is initiated by the addition of monovalent cations (Na, K), with divalent cations (Mg, Ca, Zn) mediating the compaction, bundling, crosslinking, and aggregation of vimentin filament networks.^23–25^ Additionally, the single cysteine residue (C328) located in the rod domain of human vimentin is considered a redox sensor. Interactions between C328 and transition metal ions, such as Zn(II), affect filament response to electrophilic and oxidative stress, as well as vimentin polymerization and filament reorganization.^26–30^ Vimentin also interacts with copper (Cu) and influences Cu-promoted migration of adipose-derived stem cells^31^ and critical Cu-dependent functions in the regression of cardiac cell hypertrophy.^32^ Yet, the molecular basis for the metal-induced structural changes and functions of vimentin are still unclear.

In comparison to the head and rod domains, the conformational landscape of the vimentin tail domain is underexplored. The tail domain is highly dynamic and flexible, containing conserved motifs including a helix-terminating motif and beta-turn motif, but also contains some regions with ordered structure that undergo conformational transitions during filament assembly and higher-order organization.^33^ Vimentin filaments have been described as resembling a “bottlebrush” structure, where the tails extend out from the filament core (Figure 1B).^34^ This disposition of the tails was further supported by immunofluorescence imaging demonstrating the appearance of a “cloud” of tails around the core of vimentin IFs.^35^ Surface exposed tail domains likely facilitate tail-dependent modulation of filament structure and filament-filament interactions. Although once believed to serve nonessential functions, the vimentin tail domain is crucial for the distribution of vimentin inside of cells,^36^ mediating the direct binding and interactions with other proteins,^37^ and regulating the structural and mechanical properties of filament networks through interactions with divalent metal ions.^23^ Deletion of the last 11 residues (NETSQHHDDLE) at the C-terminus disrupts metal-responsive processes and alters filament organization,^24^ suggesting that the tail domain may contain a direct metal binding site, the presence of which could drive metal ion alterations of vimentin IF structure. A complete understanding of the mechanisms that govern the metal ion interactions of the tail domain and how these interactions influence vimentin structure and dynamics necessitates an in-depth characterization of both the conformational and direct metal binding properties of the tail region.

In this work, we implemented a bipartite approach involving parallel investigations of peptide and protein tail domain fragments, to elucidate and characterize the specific protein-metal interactions of the tail domain. Analysis by mass spectrometry and spectrophotometry reveal direct metal interactions. Nuclear magnetic resonance (NMR) measurements characterize the conformation of the entire isolated vimentin tail domain. The structural effects of Cu(II) on the tail domain fragments were probed by circular dichroism and NMR techniques. Our findings point to the tail domain serving as a key metal binding region of vimentin and provide further support of the important interplay between metal ions and vimentin IF physiology and pathology.

## Results

### Preparation of tail domain fragments

To investigate the specific interactions between the tail domain of vimentin and metal ions, we produced vimentin C-terminal tail domain peptide (VimTPep) and protein (VimT) fragments. Figure 1A shows the three domains of the full-length vimentin protein and highlights the C-terminal regions of the protein that were examined in this study. The tail domain (VimT) spans residues E405–E466 (62 amino acids), with the 11-amino acid long peptide (VimTPep) corresponding to residues N456–E466. VimTPep was prepared using Fmoc-based solid-phase peptide synthesis including an N-terminal acetylation step and purified by reverse phase-high performance liquid chromatography (RP-HPLC). The VimT protein fragment was recombinantly expressed in *E. coli*, purified by fast performance liquid chromatography (FPLC), and isolated using anion exchange (AEX) chromatography (see Experimental procedures section for full preparation details). VimTPep, which contains the final 11 residues of the vimentin tail domain (NETSQHHDDLE) reported to be crucial for divalent cation interactions,^24^ was used as a model to initially screen for direct metal binding. To evaluate the potential role of the 11 residues, as well as the other regions of the tail domain in direct metal ion interactions, parallel and systematic peptide and protein structural and metal binding characterization studies were performed.

### Direct metal binding to vimentin tail domain peptide (VimTPep)

Previous work has shown that tail-truncated vimentin proteins, particularly deletion of the last 11 residues (N456–E466), are deficient in Ca(II) and Mg(II)-mediated crosslinking of filament networks.^24^ We thus sought to determine if metal ions could directly interact with VimTPep. To assess the direct metal ion interactions, VimTPep was incubated with different metal salts and the resulting adducts were analyzed by electrospray ionization mass spectrometry (ESI-MS). Initial ESI-MS experiments were carried out on a single quadrupole LC/MS instrument and analysis of non-acetylated VimTPep with various metals resulting in the emergence of peptide/metal adducts only in the presence of an equimolar amount of Cu(II) (Figure S2). These studies were followed up with high-resolution ESI-MS (Thermofisher LTQ XL Orbitrap) of the N-acetylated forms of VimTPep to further probe species that are in low abundance or have low ionization potential. The mass spectra of apo VimTPep show two predominant peaks corresponding to singly and doubly charged molecular ions with *m/z* values of 1366.55 ([M + H]^+^) and 683.78 ([M + 2H]^2+^), respectively (Figure 2A). The apo VimTPep spectra also exhibit potassium bound VimTPep adducts corresponding to [M + K]^+^ (*m/z* = 1401.51), [M + K]^2+^ (*m/z* = 702.75), and [M + 2H + K]^2+^ (*m/z* = 721.73) (Figure 2A). The addition of Cu(II) at a 1:1 peptide-to-metal ratio results in the presence of a new doubly charged mass peak with an *m/z* value of 714.23 ([M + Cu]^+^), indicating the formation of a Cu(II)/VimTPep adduct (Figure 2B). Similarly, the addition of 1 molar equivalent of Zn(II) results in the presence of a Zn(II)/VimTPep adduct with an *m/z* value of 714.73 ([M + Zn]^+^) (Figure 2C). The peptide/metal adducts are further confirmed by the observation of distinct Cu and Zn isotopic distribution patterns (Figure S3). The formation of peptide/metal adducts were not observed with the addition of 1 equivalent of Ca(II) and Mg(II) (Figure S4). Analysis by ESI-MS therefore suggests that both Cu(II) and Zn(II) can directly bind to VimTPep at a 1:1 peptide-to-metal ratio, forming a complex that persists during ionization and in the gas phase required for ESI-MS analysis.

**Figure 2.**
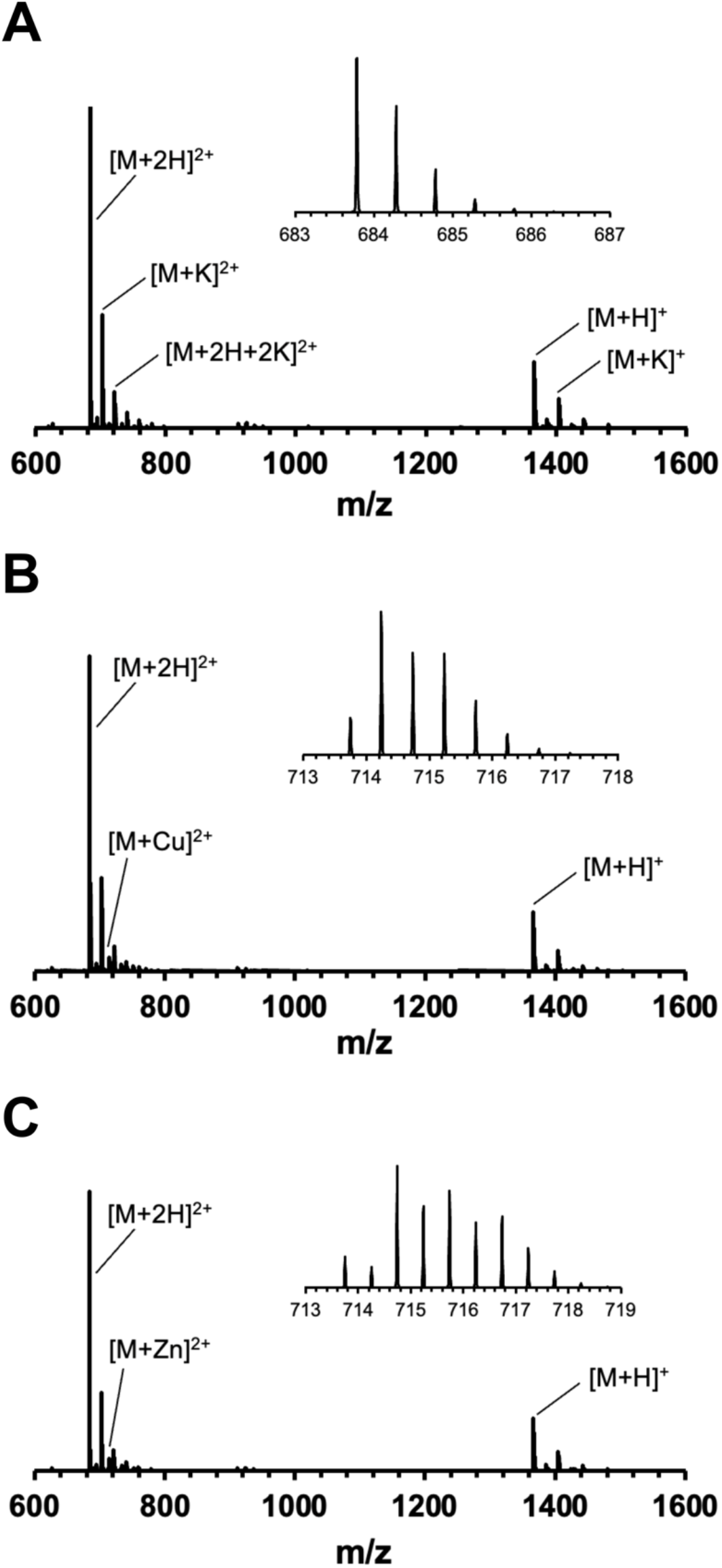
Metal binding to VimTPep was screened by ESI-MS. The spectrum of (A) apo VimTPep (50 µM) exhibits two predominant mass peaks corresponding to [M+H]^+^ and [M+2H]^2+^ molecular ions. The addition of equimolar (B) Cu(II) or (C) Zn(II) to VimTPep solutions results in the presence of new peaks indicative of the formation of VimTPep/Cu(II) and VimTPep/Zn(II) adducts, at a 1:1 peptide-to-metal ratio. The presence of the peptide/metal adducts demonstrates that both copper and zinc divalent cations may directly interact with VimTPep in the gas phase.

### The binding of Cu(II) to VimTPep can be characterized using a chromophoric chelator

Having observed the potential of Cu(II) and Zn(II) to bind to VimTPep, the Cu(II) and Zn(II) interactions of VimTPep were further characterized by electronic absorption spectroscopy to approximate the binding affinities in aqueous solutions. As the Cu(II) d-d band has a low molar extinction coefficient and Zn(II) is spectroscopically silent, peptide/metal interactions were evaluated by challenging metal-bound chromophoric chelators with VimTPep; in this way, the known affinities between metal and chromophoric chelators could be applied to approximate and compare VimTPep-metal binding affinites. Additionally, these competitive metal binding assays enable the use of low peptide and protein concentrations. Two well-characterized chelators, 1,10-phenthroline (phen) and zincon (ZI) were used to approximate Cu(II) and Zn(II) binding affinities, respectively.^25,26^

Solutions of VimTPep in 20 mM phosphate buffer (pH 7.4) were prepared and titrated into either a buffered solution of 40 µM phen and 10 µM Cu(II) or 20 µM ZI and 10 µM Zn(II), and the data was evaluated at a range of 10–200 µM peptide. The Cu(II)-phen complex exhibits a characteristic charge-transfer band at λ = 265 nm^26^ that can be monitored and used to assess the competition between tail domain fragments and phen. The titration of VimTPep into a solution containing a complex of Cu(II)-phen resulted in a decrease in the absorption of the charge-transfer band, indicative of the competitive binding of Cu(II) to the tail domain peptide (Figures 3A). Using the data from the titration experiment and following methods outlined previously^38–40^ the binding affinity of Cu(II) to VimTPep is estimated to be between log*K* 7.5 and 8.7, or a maximum *K*_d,app_ value of 32 nM (Figure 3A and Table S1).

**Figure 3.**
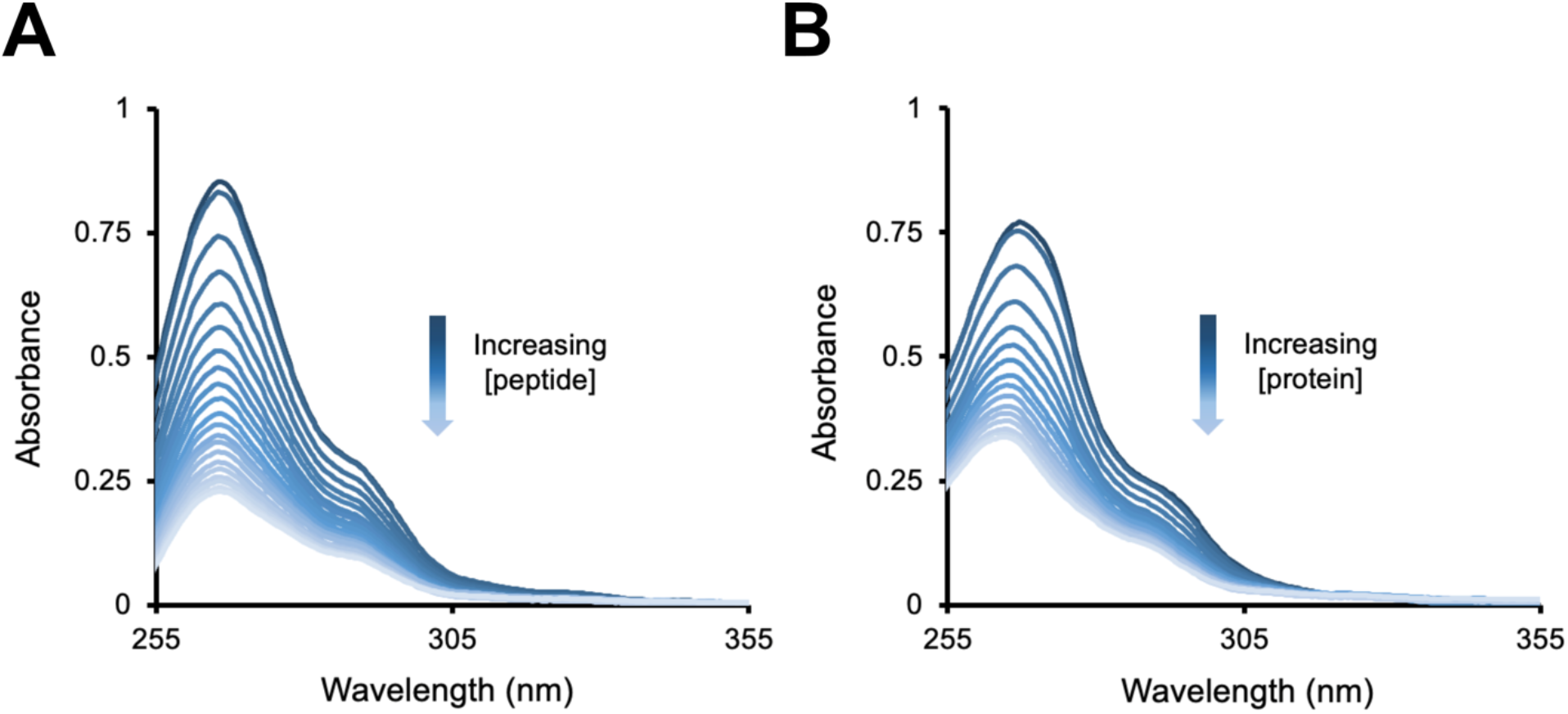
Assessment of the Cu(II) binding affinity to vimentin tail domain fragments using the chromophoric ligand, 1,10-phenanthroline. (A) Electronic absorption spectra of 10–200 µM peptide titrated into a solution of 10 µM Cu(II) and 40 µM phen in 20 mM phosphate buffer (pH 7.4). The competitive binding of VimTPep to Cu(II) was determined to have an approximate binding affinity of log *K* between 7.5– 8.7 (or K_d,app_ = 2–32 nM). (B) Electronic absorption of 6–100 µM VimT protein titrated into a solution of 10 µM Cu(II) and 40 µM phen in 20 mM phosphate buffer (pH 7.4). The same approach used for VimTPep was extended to the VimT fragment to determine the competitive binding of VimT to Cu(II). The estimated binding affinity of Cu(II) to VimT is log *K* between 7.5–8.6 (or K_d,app_ = 2–27 nM).

As the Zn(II)/VimTPep complex was also observed in MS studies, the chromophoric ligand ZI was applied to characterize Zn(II) binding. However, initial work with the ZI assay using manually synthesized vimentin tail domain peptides prepared in buffered solutions, proved to be unsuccessful resulting in either inconclusive data or ineffective competitive binding (data not included). From these observations, it is likely that Zn(II) binds weakly to VimTPep or that it does not bind in a competitive manner that can be assessed using the ZI-Zn(II) complex. Based on the results gathered using ZI and phen as competitive reporters of metal binding to VimTPep, subsequent experiments focused on examining the Cu(II) binding of the tail domain and investigating the basis of these metal-protein interactions.

### VimTPep and the vimentin tail domain fragment (VimT) have similar Cu(II) binding affinities

The same approach was used to characterize Cu(II) binding to the larger tail domain fragment, VimT, to determine the influence of the N-terminal tail domain region on the nanomolar affinity of VimTPep for Cu(II). Solutions of VimT were prepared in 20 mM phosphate buffer (pH 7.4) and titrated into a buffered solution of 40 µM phen and 10 µM Cu(II), to evaluate a concentration range of 10–200 µM protein. Similarly to VimTPep, the titration of VimT into Cu(II)-phen solutions led to a decrease in the absorption of the charge-transfer band, demonstrating the competitive binding of Cu(II) to the tail domain protein fragment (Figure 3B). With this method, the approximate binding affinity of Cu(II) to VimT is estimated to be log*K* between 7.5–8.6 or K_d,app_ = 2–27 nM (Figure 3B and Table S2), which is markedly similar to VimTPep. This result suggests that VimTPep and VimT share common Cu(II) binding motifs.

### Cu(II) binding alters the conformation of VimTPep and VimT

The structural consequences of Cu(II) binding to VimTPep and VimT were examined by circular dichroism (CD) spectroscopy. Both fragments are largely unstructured as indicated by a characteristic minimum around 198 nm in the far-UV CD spectra of VimTPep (Figure 4A) and VimT (Figure 4B). The addition of 0–10 equivalents of Cu(II) to VimTPep shows a decrease in the random coil conformation of the peptide, as demonstrated by reduction in the ellipticity of the negative band between 190–200 nm (Figure S5), with the most significant changes occurring between 0–1 molar equivalents of Cu(II). This range was further probed with smaller titration increments (Figure 4A), revealing two major changes: a decrease in the negative ellipticity values at 198 nm (Figure 4A, orange arrow, and Figure S6A) indicative of the loss of random coil conformation, and an increase in the negative ellipticity at 224 nm (Figure 4, red arrow, and Figure S6B). However, the observed changes in the VimTPep spectra between 210–230 nm cannot be attributed to any Cu(II)-induced secondary structure (Table S3).

**Figure 4.**
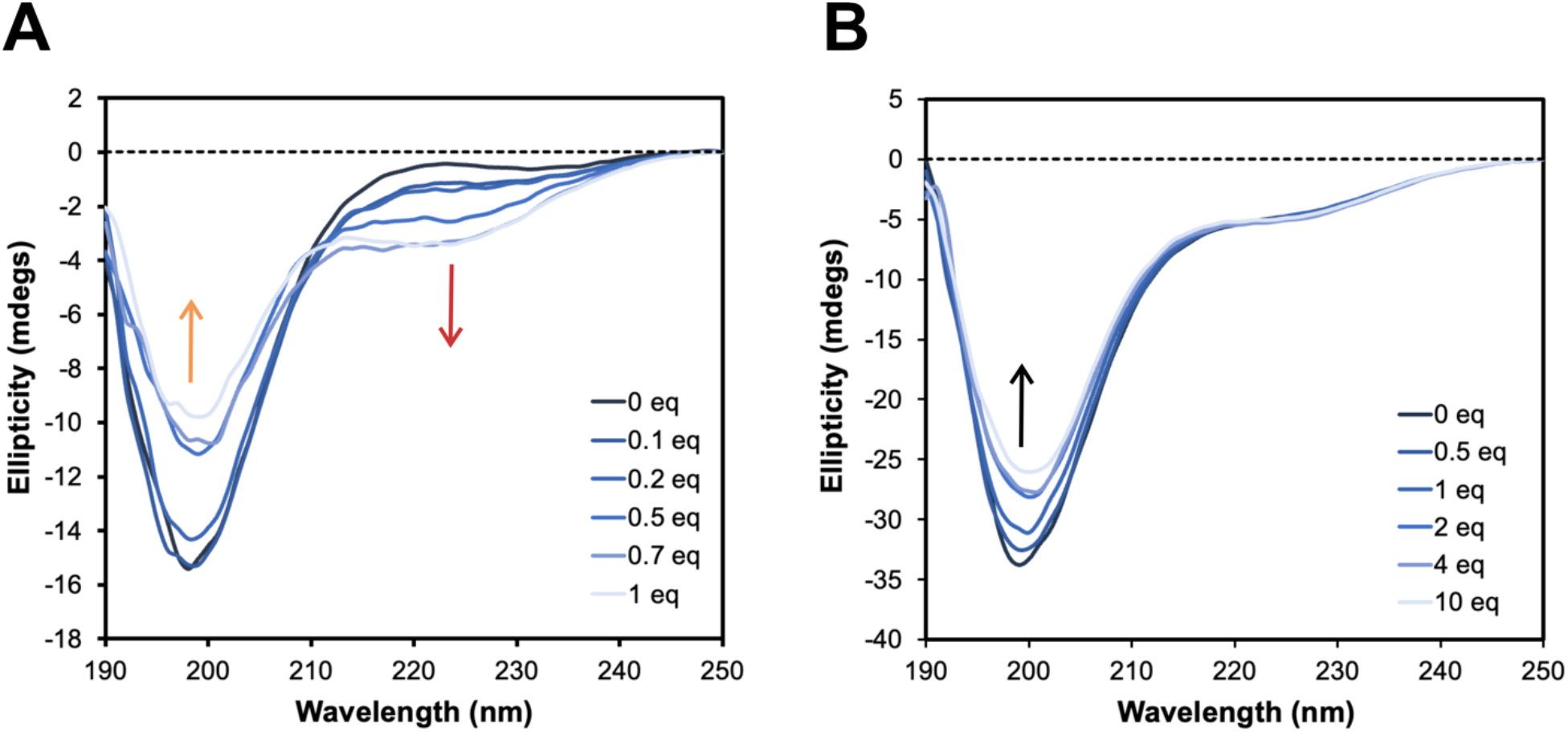
Circular dichroism spectra of VimTPep (100 μM) and VimT (∼37 μM) in 20 mM phosphate buffer (pH 7.4). (A) CD spectra showing changes in the random coil conformation of VimTPep upon the gradual addition of 0 to 1 Cu(II) molar equivalents (eq), demonstrated by the reduction of ellipticity at 198 nm (orange arrow). The CD spectra also exhibit an increase in negative ellipticity values at 224 nm (red arrow). The addition of 0 to 1 molar equivalents of Cu(II) corresponds to 0 to 100 μM Cu(II). (B) CD spectra showing a modest reduction in the random coil conformation of VimT upon the addition of 0 to 10 eq of Cu(II), shown by the slight reduction of ellipticity at 198 nm (black arrow). No significant changes in ellipticity are observed from 210–230 nm. The addition of 0 to 10 eq of Cu(II) to VimT corresponds to 0 to 370 μM Cu(II).

A similar Cu(II) titration was used to probe changes in the secondary structure of VimT. VimT concentrations of at least 14 μM were required to properly assess the Cu(II) interactions, with optimal CD spectra obtained with VimT concentrations of ∼37 μM (∼0.3 mg/mL) (Figures S7 and 4B). CD analysis of VimT demonstrated a decrease in random coil conformation upon the addition of 0–10 or 0–0.5 equivalents of Cu(II) at both pH 7.4 (Figure 4B) and pH 6.5 (Figure S8) (the pH used for NMR experiments), respectively. In contrast to VimTPep, the titration of Cu(II) results in a smaller magnitude decrease in ellipticity at 198 nm for VimT, with no observed changes between 210 and 230 nm (Figure 4B). The addition of Cu(II) to VimT results in more subtle changes in secondary structure compared to VimTPep (Figure S6C), likely attributed to the presence of disordered segments that remain unaffected within VimT that strongly contribute to the average CD signal. CD analysis of the data from Figure 4 using the program BeStSel^41,42^, predicted a disordered classification for both VimTPep and VimT.

### VimT conformation is largely disordered and dynamic

Nuclear magnetic resonance spectroscopy (NMR) was used to further probe the structural changes of VimT occurring upon interaction with Cu(II). As the isolated tail domain of vimentin has not been previously characterized by solution NMR, we first analyzed the tail domain protein fragment in the absence of metals. The ^1^H–^15^N heteronuclear single quantum coherence (HSQC) spectrum of uniformly ^13^C, ^15^N-labeled VimT shown in Figure 5A exhibits the narrow ^1^H dispersion expected for a largely disordered protein, corroborating the structural properties observed by CD at 0 Cu(II) equivalents. To determine residue-specific structural information, a standard set of 3D experiments were recorded on the uniformly ^13^C, ^15^N-labeled VimT sample for the purposes of sequence-specific resonance assignment. Signals were observed for 62 non-proline residues, out of which 51 (or 82%) could be unambiguously assigned in the 67-residue VimT fragment used in our experiments, which contains 4 proline residues. The assignments are overlaid in the ^1^H–^15^N HSQC spectrum in Figure 5A. Of the unassigned signals, 4 were barely above the noise level. All experimental and data acquisition parameters are provided in Table S4. The assigned NMR chemical shifts have been deposited in the BMRB (accession number 52964).

**Figure 5.**
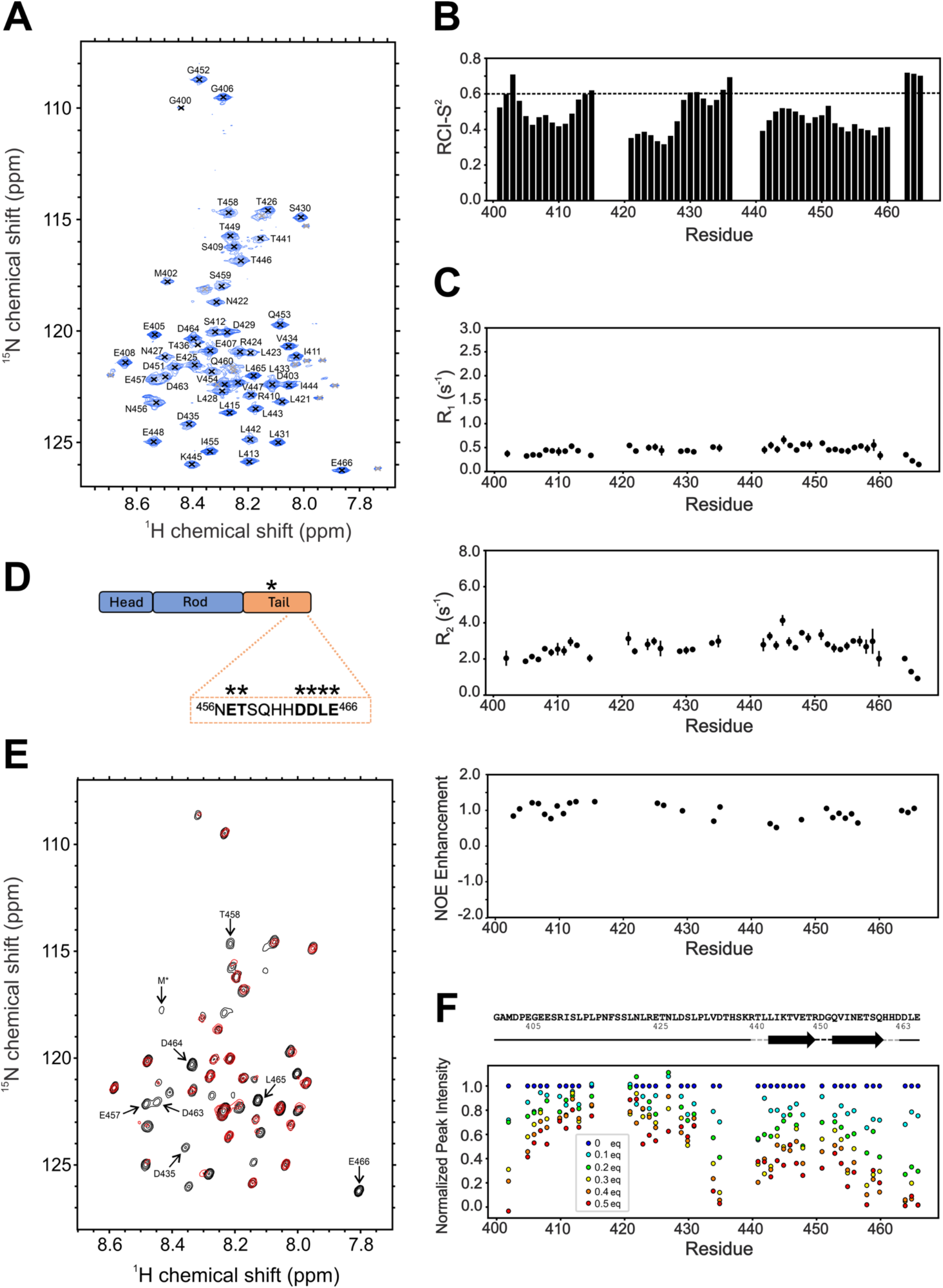
NMR characterization of vimentin tail domain protein fragment in the absence and presence of Cu(II). (A) ^1^H–^15^N heteronuclear single quantum coherence (HSQC) spectrum of ^13^C, ^15^N-labeled VimT protein in 20 mM phosphate buffer (pH 6.5), recorded at 600 MHz ^1^H frequency. Each peak corresponds to an amide proton-nitrogen pair for a single non-prolyl residue in the protein sequence, serving as a protein “fingerprint”. The distribution and location of peaks indicate that VimT is primarily disordered under these measurement conditions. Unambiguously assigned signals, 51 in total, are labeled in black and low signal-to-noise unassigned peaks are indicated with grey X-marks. (B) TALOS-N ‘random coil index’ order parameter (RCI-S^2^) classification of backbone rigidity based on the assigned VimT NMR chemical shifts. Residues with values of RCI-S^2^ ≤0.6 are indicative of dynamic regions.^45^ (C) The residue-specific measured R_1_ relaxation rates, calculated R_2_ relaxation rates from R_1_ and R_1ρ_ measurements, and ^1^H–^15^N heteronuclear NOE enhancements of VimT. All relaxation data were recorded at 600 MHz ^1^H frequency. (D) Diagram of the regions within VimT that are most affected by the addition of Cu(II) indicated by an asterisk, with a zoomed in view of the specific residues (in bold) within the last C-terminal 11-amino acids that experienced the greatest signal loss. (E) ^1^H–^15^N HSQC spectral overlay of apo protein (black) and 0.5 equivalent (red) of Cu(II) at pH 6.5, recorded at 800 MHz ^1^H frequency. Arrows point to the signals that are most quenched by the paramagnetic Cu(II). M^*^ corresponds to the methionine residue from the 6xHis purification tag that remains after cleavage by the TEV protease. (F) Residue-specific integrated peak intensities for each Cu(II) equivalent (eq) normalized to the 0 eq Cu(II) spectrum. The cartoon above the data points in panel (F) shows the Alpha Fold 2.0 secondary structure predictions of the vimentin monomer (Figure S9), with black arrows and dashed line indicating high confidence beta-strand and turn predictions. Gray dashed lines represent low confidence beta-strand predictions and the solid black lines indicate regions with no confident structure prediction.

Further structural information of the tail domain was obtained by using the protein backbone structure prediction program TALOS-N^43,44^ and from NMR relaxation experiments. TALOS-N analysis of the assigned VimT chemical shifts resulted in predominantly random coil predictions, with most residues having a random coil index order parameter (RCI-S^2^)^45,46^ value less than or equal to 0.6 (Figure 5B), reflecting the largely disordered nature of the tail domain. Residues 400–403, 416–420, 436–440, 461–462, and 466 are missing RCI-S^2^ values due to missing NMR chemical shift assignments or TALOS-N predictions. NMR relaxation measurements were used to more precisely probe the residue-specific global and local motions within VimT (Figure 5C). Spectra reporting on the longitudinal relaxation rates (R_1_), rotating frame longitudinal relaxation rates (R_1ρ_), and heteronuclear ^1^H–^15^N nuclear Overhauser effects (NOE) were recorded. The transverse relaxation rates (R_2_) were calculated based on the measured R_1_ and R_1ρ_ values, following approaches previously reported.^47,48^ The residue-specific R_1_ and R_2_ rates of VimT are consistently small across the sequence (Figure 5C) and there were no areas of large NOE enhancement values (Figure 5C). Taken together, the lack of any significant deviation from values expected for an intrinsically disordered structure or evidence for conformational exchange in the data, suggests that VimT is primarily disordered in the absence of Cu(II).

### Cu(II) binds to last 11 amino acid segment and center of VimT with minimal changes in structure

To determine the residues mediating the binding of Cu(II) to VimT, a range of Cu(II) molar equivalents (0, 0.1, 0.2, 0.3, 0.4, 0.5) were titrated into a solution of ^13^C, ^15^N-labeled VimT and a series of ^1^H–^15^N HSQC spectra were recorded (Figure S10). Paramagnetic Cu(II) quenches the NMR signal from residues in close proximity to the Cu(II). Figure 5E shows that the addition of 0.5 equivalents of Cu(II) drastically reduces the signal intensity of a subset of residues in VimT. The integrated peak intensities shown in Figure 5F reveal that residues after L433 are significantly more affected by the presence of Cu(II) than the N-terminal half of VimT. Consistent with the expected divalent metal binding role of residues N456–E466, these sites exhibit Cu(II)-dependent signal loss (Figures 5D–F). However, residues L433–T436 also undergo strong signal loss in these spectra, supporting a role for other regions of VimT in metal binding. The weak methionine (M^*^) signal, which is an unremoved residue of the purification His-tag, appears to disappear upon the addition of Cu(II), but a decrease in nearby signals is not observed suggesting that this site is not a major contributor to Cu(II) binding (Figures 5E and 5F). Note that residues H437–R440 are not observed in any of our NMR spectra, preventing analysis of Cu(II)-dependent effects in this region. Changes in peak position with the introduction of Cu(II) were not observed, for all signals, regardless of the extent of the quenching. Residues relatively unaffected by the quenching, mainly those preceding L433, therefore remain disordered irrespective of Cu(II) binding. For sites exhibiting a severe reduction in signal, the remaining intensity arises from VimT molecules unbound to Cu(II) and still in the disordered conformation, consistent with sub-stoichiometric Cu(II) concentrations used in the titration. VimT molecules directly bound to Cu(II) have unmeasurable NMR signals due to the paramagnetic broadening effect. Overall, the unchanged positions for residues not directly involved in the binding suggest that the conformation of the N-terminal portion of VimT remains mostly unaffected upon Cu(II) binding, which is consistent with the findings obtained by CD. Finally, the lack of strong signals for residues H437–R440 and H461–H462 suggests that these sites might be in a conformational exchange with a Cu(II)-binding competent conformation in the absence of Cu(II).

## Discussion

In this study, we performed an in-depth analysis of the structural and metal binding properties of the vimentin tail domain and found that Cu(II) directly binds to the domain. From mass spectrometry analyses, the divalent cations Cu(II) and Zn(II), but not Mg(II) and Ca(II), bind tightly enough to VimTPep for detection in the gas phase. Using a bipartite approach involving parallel investigations of the tail domain peptide (VimTPep, residues N456–E466) and tail domain protein fragment (VimT, residues E405–E466), we have shown that the last 11 residues (NETSQHHDDLE) of the C-terminus are likely the primary site responsible for mediating Cu(II) interactions. Additional residues in the center of the tail domain were also found to play a role in Cu(II) binding. In the absence of Cu(II), VimT is primarily disordered. The precise conformation of the Cu(II)-bound VimT domain was not detectable in our NMR measurements due to the paramagnetic nature of Cu(II). Intriguingly, Cu(II) has minimal effects on the N-terminal portion (residues E405–S430) of the tail domain, which is disordered regardless of Cu(II)-binding.

While the structure of the rod domain of vimentin has been extensively investigated, less is known about the configuration of the tail domain. In the context of the full-length vimentin filaments, it has been proposed that the tails protrude from the filament core forming a “bottlebrush” like structure, enabling vimentin tails to adopt various conformations and regulate filament organization and interactions.^49^ Recently, cryo-electron microscopy (cryo-EM) analysis of *in vitro* prepared vimentin filaments showed the tail domain adopting structure, although the resolution for the domain was not sufficient to provide a specific conformation or to conclude the entire domain was structured.^18^ In agreement, an EPR investigation of vimentin filaments assembled *in vitro* showed a portion of the tail domain (residues K445–G452) in close proximity to tail domains from other vimentin monomers, possibly due to the formation of structure.^33^ Also consistent with the EPR result, the AlphaFold 2.0^50^ prediction for the vimentin monomer (Figure S9) weakly predicts two beta-strand structures folded into a hairpin structure for residues L443–Q460. In the absence of metal ions, our CD, NMR chemical shift measurements, and NMR relaxation measurements of VimT (Figures 4B, 5B, and 5C respectively) show that the tail domain in isolation is primarily disordered in solution, which is not surprising since our VimT is likely monomeric in solution. Disorder in the N-terminal segment prior to residue L433 is consistent with both the AlphaFold 2.0 prediction and the EPR study. Comparison to the cryo-EM results is not possible due to the lack of high resolution density for the tail domain in the reconstructed electron density.

Previous work has shown that metal ions such as Mg(II) and Ca(II) can interact with the vimentin tail domain and modulate the structure of vimentin filament networks.^24^ In the presence of Mg(II), Ca(II), Cu(II), and Zn(II) ions, we were able to observe and assess only Cu(II)-tail domain interactions. Our mass spectrometry results (Figure 2B) indicate that Cu(II) binding can occur at a 1:1 ratio of metal-to-tail domain but does not preclude the binding of other divalent metal ions at higher ratios or in a manner dependent on the head and rod domains of vimentin. Detailed characterization confirmed that both VimTPep and VimT fragments can bind directly to Cu(II) (Figures 4 and 5). Similar approximate nM Cu(II) binding affinities were measured for both fragments, suggesting common Cu(II) binding sites. CD analysis shows that Cu(II) induces changes in the random coil conformation of VimTPep, but the influence is unclear for VimT as the CD spectra for that fragment are likely dominated by disordered segments of the tail domain that do not exhibit structural differences upon binding to Cu(II).

Vimentin is considered to be a Cu-binding protein,^32^ but its exact Cu binding sites are unknown. Solution NMR studies (Figures 5E and S10) of VimT enabled us to obtain greater detail of the Cu(II)-tail domain interactions. A series of ^1^H–^15^N HSQC measurements at increasing Cu(II) concentrations reveals the quenching of residue-specific signals, with the greatest loss in signal intensity primarily exhibited by residues between N456–E466, corresponding to the 11 residues of the VimTPep fragment. In this range, signals for 6 of the 11 amino acids (E457, T458, D463, D464, L465, E466) are most affected by Cu(II).

Residue D435, located in the center of the tail domain, also experiences a large reduction in signal intensity. When inspecting the primary sequence of VimTPep, we expected to also observe interactions between Cu(II) and the two histidines (H461 and H462) present in the sequence. However, the missing NMR signals for residues H461–H462 prevented analysis of the binding of Cu(II) to these sites. The lack of NMR signal for these sites in the absence of Cu(II) could be due to signal broadening arising from the conformational exchange between the bound and unbound forms of VimT or simply the formation of a less disordered structure. In general, the distal residues of the C-terminus (after L433) underwent the greatest changes in signal after the addition of Cu(II), compared to the residues proximal to the N-terminus of the tail domain. Moreover, paramagnetic Cu(II) dictates that the residues coordinating the metal are not observable in our NMR experiments. Nevertheless, the lack of new signals or shifted positions of visible peaks, indicates that Cu(II) does not induce significant structure in the segments of the tail domain not participating in the binding interaction, consistent with the small changes observed in the CD spectra of VimT in the presence of Cu(II).

Our NMR findings are supported by AlphaFold 2.0^50^ (AF2) and 3.0^51^ (AF3) predictions of the structural and metal binding properties of the vimentin tail domain. AF2 predicts with high confidence beta-strand and turn regions between residues L443–Q460 (see Figures 5F and S9). NMR signals are mostly unobserved for residues at the ends of the predicted beta-strands and turn region (H437–R440, R450, and H461–H462), consistent with increased order and reduced molecular motions at these sites. In addition, the AF3 model of VimT with Cu(II) shows residues near the beginning and end of the predicted beta-strand region (H437, H461, D463) to coordinate Cu(II) (Figure S11B), in agreement with the results of our NMR titration. AF3 models of higher-order oligomers of vimentin (e.g., dimers and tetramers, models not shown) with the addition of Cu(II) resulted in unreliable predictions and do not predict the expected rod domain structures. The full-length vimentin monomer model predicts the binding of Cu(II) to C328 rather than the tail domain (Figure S11D), which is expected given previous reports of the involvement of C328 residue in mediating the binding of divalent metal ions.^28^ Therefore, the AF3 model of the VimT fragment with Cu(II) reveals additional Cu(II) binding regions in the tail domain that are not reflected in the full-length AF3 vimentin monomer model.

Vimentin and other IF proteins are subject to several post-translational modifications (PTMs) that fine-tune their structure and function.^52,53^ Phosphorylation is the most significant and well-studied IF PTM. Vimentin contains multiple phosphorylation sites in both the head and tail domains that result in the reorganization of mature filament structure, regulating key functions like cell motility and division.^54–57^ Mutations or changes in the phosphorylation sites of vimentin have been linked to disease pathogenesis.^58–61^ Tail domain residues T457 and S458 have been identified as sites for phosphorylation^62,63^ which are within the VimTPep segment investigated in our study and were found to interact with Cu(II). It is known that the electrostatic properties of IF tails are altered by phosphorylation,^34^ but the relationship between vimentin tails, metal ions, and PTMs, has not been delineated. The colocalization of phosphorylation sites with metal binding regions suggest that the tail domain is a vital component in the mechanisms regulating vimentin IF function and dysfunction.

### Concluding Remarks

The architecture of vimentin IFs is remarkably diverse and complex, which is demonstrated by the constant remodeling of vimentin filaments in response to various cellular conditions and stimuli.^64^ Here, we implemented a two-part, peptide and recombinant protein fragment approach to help unveil the characteristics of the vimentin tail domain. Our data strongly supports the ability of the tail domain, in isolation, to mediate direct interactions with Cu(II). Earlier work demonstrated that the interaction between divalent cations and the last 11 C-terminal tail domain residues is important for effective crosslinking of vimentin filaments.^24^ Our findings provide further evidence that this segment of 11 amino acids in the tail domain is a major site for the binding of metals. This work presents a comprehensive experimental framework for characterizing domain-specific IF protein-metal interactions and provides a deeper understanding of the molecular mechanisms that influence the macroscopic physiological and pathological properties of IFs.

## Experimental procedures

### Peptide synthesis and purification

Wild-type C-terminal tail domain vimentin peptide (NETSQHHDDLE) (VimTPep) was synthesized manually or using a CEM Liberty Blue 2.0 automated microwave peptide synthesizer following standard Fmoc solid-phase peptide synthesis (SPPS) strategies. VimTPep generated by manual SPPS had no N-terminal modifications, while peptides made by automated SPPS were N-acetylated. The acetylated form of the peptides (Ac-NETSQHHDDLE) was produced to best represent the native state of the peptide within the larger protein environment using acetic anhydride. Peptides were synthesized at 0.1–0.2 mmol scales using Wang resin preloaded with Fmoc-Glu(OtBu)-OH. Synthesis began with a 20 min resin swelling step. Amino acid couplings were performed using CEM-optimized microwave synthesis conditions: DIC and Oxyma Pure activating reagents, 1 min deprotection at 90 °C, followed by a 2 min coupling at 90 °C. After the coupling of the last amino acid, the final step of the synthesis was the deprotection of the N-terminal residue. Upon synthesis completion, all resin/peptide material was removed from the peptide synthesizer and washed with DMF (5x), followed by DCM (5x), and dried for 1–2 hours at room temperature. The peptide and protecting groups were cleaved from resin using a 10 mL solution containing a 95:2.5:2.5 ratio of TFA/water/TIPS. The resin in solution was allowed to shake at room temperature at 145 rpm for 3–4 hours. The cleavage solution was slowly transferred into conical tubes containing 40 mL of chilled diethyl ether to precipitate the peptide material, centrifuged for 10 min at 3,900 rpm, and the supernatant solution decanted. An additional 40 mL of chilled diethyl ether was poured into the tubes to wash the crude peptide pellet and centrifuged for an additional 10 min at 3,900 rpm, followed by decanting the supernatant. These steps were performed a total of three times. After the final decanting, the resulting crude peptide material was dried overnight in a vacufuge and stored at -20 °C until purification.

The peptide was purified by reverse phase-HPLC using an Agilent Technologies 1260 Infinity II HPLC instrument equipped with an Infinity II UV-Vis detection system. The crude VimTPep was purified using an Atlantis T3 Prep OBD C18 column (100 Å, 5 µm, 19 × 250 mm) set to a flow rate of 17.6 mL/min with a gradient of solvent A (water with 0.1% FA) and solvent B (acetonitrile with 0.1% FA) as follows: 5% solvent B held for 5 min, linear increase to 45% of solvent B from 5-85 min, and final increase to 100% of solvent B from 85-90 min. VimTPep eluted between 20-25 min or at a ratio of 90-87.5% A/10-12.5% B. The purified peptide was confirmed by electrospray ionization (ESI) or matrix-assisted laser desorption ionization (MALDI) mass spectrometry, lyophilized, and kept at -20 °C until use.

### Protein production and purification

Human wild-type vimentin tail domain protein (VimT), in a pHIS-parallel^65^ plasmid containing residues E405–E466 and an N-terminal TEV-cleavable His-tag, was recombinantly expressed in BL21(DE3) *E. coli* cells. All bacterial growth was performed in a shaker incubator set to 37 °C and 220 rpm. For all VimT protein (unlabeled and labeled), bacterial cells were grown in 2 L of Luria broth media, with 100 μg/mL ampicillin, to an optical density at 600 nm (OD_600_) between 0.6–0.8. Protein growth was induced with the addition of 0.5 mM of isopropyl β-D-thiogalactopyranoside (IPTG). The bacterial cells were grown for an additional 3 h and harvested by centrifugation at 6,000 g for 10 min. The resulting cell pellets were flash frozen in liquid nitrogen and stored at -80 °C until purification. For ^13^C and ^15^N isotopically labeled VimT protein, the cells from 2 L of Luria broth media at an OD_600_ between 0.6–0.8 were harvested by centrifugation at 6,000 g for 10 min and then transferred into 1 L of M9 minimal media containing 100 μg/mL ampicillin, 2.0 g of U-^13^C_6_ D-glucose, and 1.0 g of ^15^N ammonium chloride. The culture was incubated with shaking at 37 °C and 220 rpm for 30 min, then protein production was induced with 0.5 mM IPTG. The bacterial cells were grown for an additional 3 h and harvested for a second time by centrifugation at 6,000 g for 10 min. The final cell pellets were flash frozen in liquid nitrogen and stored at -80 °C until purification.

All VimT protein was IMAC affinity purified using a Bio-Rad NGC Discover 10 FPLC system and followed the same steps. The cell pellets (∼1–2 g wet cell mass) were thawed on ice and resuspended in a solution of 6M guanidinium hydrochloride, 1% v/v Triton X-100, 50 mM tris(hydroxymethyl)aminomethane (Tris-HCl) pH 7.5, 500 mM sodium chloride, 3 tablets of Mini EDTA-free Pierce Protease Inhibitor, and 0.25 mg/mL of hen egg white lysozyme. The resuspended cell solution was sonicated in an ice water bath using a Branson SFX-250 Sonifier equipped with a ¼ inch microtip in pulsed mode using the following settings: 0.3 s on, 3 s off, 1 min total sonication on time, at 30 % power (total time of 20 min). After sonication, the lysed cells were centrifuged at 4 °C at 75,600 g for 30 min, to separate the insoluble material. The resulting supernatant solution was flowed over a 5 mL Bio-Rad Bio-Scale Mini Nuvia IMAC column equilibrated with 6 M urea, 20 mM 4-(2-hydroxyethyl)-1-piperazineethanesulfonic acid (HEPES) pH 7.5, and 500 mM sodium chloride; washed with 6 M urea, 20 mM HEPES (pH 7.5), 500 mM sodium chloride, and 20 mM imidazole; and eluted using a 20–200 mM gradient of imidazole, along with 6 M urea, 20 mM HEPES (pH 7.5), and 500 mM sodium chloride. The elution peak was collected in 1 mL aliquots and stored at -80 °C until further isolation/preparation of protein. The purified protein material was analyzed by SDS-PAGE with Coomassie staining and protein concentrations determined using absorbance measurements at 280 nm and a calculated molar extinction coefficient of 5,960 M^-1^ cm^-1^ (ProtParam^66^).

### Protein isolation via His-tag cleavage and anion exchange chromatography

To prevent interferences with metal binding experiments, the N-terminal 6xHis-tag (Figure S1A) was removed from purified protein using the TEV protease at a concentration ratio of 1:100 protein-to-TEV (Figure S1B). Removal of the His-tag was accomplished by adding TEV to a solution of purified VimT protein (∼4–5 mL) inside dialysis tubing and by dialyzing the TEV-VimT mixture into a 250 mL buffer solution containing 20 mM Tris-HCl (pH 7.5), 1 mM DTT, and 1 mM EDTA. The buffer solution was replaced with fresh buffer after 3 h and the dialysis continued overnight (∼19 h) at room temperature. Immediately following TEV cleavage, the VimT protein was separated from the cleavage products by anion exchange chromatography (AEX) using a 5 mL Bio-Rad EconoFit Macro-Prep DEAE column.

AEX was performed manually by attaching the DEAE column on a ring stand and running sodium chloride buffered solutions through the column. All applications to the column were done using a 10 mL syringe. The contents of the low and high sodium chloride buffers were as follows: 20 mM Tris-HCl (pH 7.5), 6 M urea, and 1 mM EDTA (pH 8) for low salt and 1 M sodium chloride, 20 mM Tris-HCl (pH 7.5), 6 M urea, and 1 mM EDTA (pH 8) for high salt. The column was first prepared following the steps outlined in the column instruction manual. In brief, 20 mL (4 cv) of ultrapure water was run through the column to remove the storage ethanol solution (20% v/v), followed by 12 mL (2.4 cv) of low salt buffer and 30 mL (6 cv) of high salt buffer, and equilibrated with 30 mL (6 cv) of low salt buffer. After column preparation, protein solution was loaded slowly onto the column and 5 mL of low salt buffer was used to assist in the complete transfer and loading of all protein material. The column was allowed to sit for 10 min after sample loading and prior to collection. A 0–300 mM sodium chloride gradient was run in 50 mM sodium chloride increments. For each step, a 10 mL solution was prepared by mixing high and low salt buffers and run over the column. Each 10 mL step was collected in two 5 mL fractions and stored at 4 °C for 1–3 days. The presence of isolated VimT protein material was confirmed in fractions corresponding to 50–100 mM sodium chloride concentrations by SDS-PAGE with Coomassie staining (Figure S1C) and the fractions stored at -80 °C until use. Since the His-tag cleaved protein material did not contain residues with an absorbance at 280 nm (0 M^-1^ cm^-1^ molar absorptivity), all protein concentrations were calculated by using absorbance measurements at 205 nm and a molar extinction coefficient of 217,310 M^-1^ cm^-1^ as detailed by Anthis and Clore^67^ and using the protein parameter calculator (accessed online at http://nickanthis.com/tools/a205.html).

### Mass spectrometry

Initial ESI-MS experiments were performed on an Agilent Technologies InfinityLab single quadrupole LC/MSD instrument. High-resolution MS experiments were carried out using a Thermofisher LTQ XL Orbitrap operated in positive ion mode at a scan resolution of 60,000 for MS scans. All VimTPep solutions at 50 or 100 μM were prepared with 5 mM ammonium acetate (pH 7.4) with an equimolar amount (50 or 100 μM) of CaCl_2_, CuSO_4_, MgCl_2_, and ZnCl_2_ metal solutions prepared in nanopure water, added to each peptide sample. Source conditions for the Orbitrap instrument were as follows: spray voltage of 3.5 kV, capillary temperature of 275 °C, and a flow rate of 0.2 mL/min. The following detection settings were used: automatic gain control target at 3.0 × 10^6^, maximum injection time of 200 ms, microscans of 1, and a scan range of 250-2000 *m/z*. Orbitrap MS spectra were analyzed using Thermo Scientific Xcalibur software, averaged over 50 scans, and exported as text files.

### UV-visible spectroscopy

All measurements were acquired using a Shimadzu UV-1900 spectrophotometer at room temperature using 1 cm path length quartz cuvettes (Starna Cells). The competitive binding studies were adapted from Stevenson et al.^39^ using the 1,10-phenanthroline (phen) ligand. The preformed complex of Cu(II)-phen was prepared by separately dissolving CuCl_2_ and phen in Milli-Q water to make 1 mM stock solutions, followed by combining Cu(II) and phen to final concentrations of 10 μM and 40 μM, respectively. The Cu(II)-phen mixture was incubated for 10 min at room temperature before the addition of peptide or protein. The VimTPep and VimT solutions were prepared using 20 mM phosphate buffer (pH 7.4) and titrated into the Cu(II)-phen at final peptide/protein concentrations ranging from 0 to 200 μM. After each peptide/protein addition the solution was equilibrated for 3–5 min at room temperature and spectra collected from 200–900 nm. The absorbance of Cu(II)-phen was monitored at 265 nm. Absorbance measurements were recorded with water as a reference followed by subtraction of buffer referenced to water, and data normalized to account for dilution. Calculations for the formation constants of Cu(II)/VimTPep and Cu(II)/VimT complexes were done following approaches previously outlined.^38–40^ The corresponding values and calculations for VimTPep and VimT are shown in Table S1 and Table S2, respectively.

### Circular dichroism spectroscopy

CD measurements were collected on a JASCO J-715 spectropolarimeter at room temperature using 1 mm path length quartz cuvettes (Science Outlet). Lyophilized VimTPep (100 μM) samples were prepared in 20 mM phosphate buffer (pH 7.4) and VimT (∼10–40 μM or ∼0.1–0.3 mg/mL) samples were prepared by dialyzing 1–4 mL of isolated protein material into 20 mM phosphate buffer at either pH 7.4 or pH 6.5. The metal stock solutions were made by dissolving metal salts in Milli-Q water at a concentration of 0.5, 1, or 10 mM. Metal solution was added to VimTPep and VimT at a range of 0–1 or 0–10 molar equivalents and each mixture was incubated for 10 min at room temperature prior to data collection. Spectral acquisition was performed by scanning 190–250 nm using a continuous scanning mode, 20 nm/min scanning speed, 2–3 averaged scans, and a bandwidth of 1.0 nm. CD spectra were baseline-corrected, buffer spectrum subtracted, and smoothed prior to analysis. The BetStSel CD spectra analysis program (accessed online at https://bestsel.elte.hu) was used to estimate the percentage of secondary structure and the “Disordered-Ordered Classification” option was applied to generally classify the disordered properties of VimTPep and VimT, based on their far-UV CD measurements.^41,42^

### NMR spectroscopy

NMR experiments were performed on Bruker 600 MHz or 800 MHz Avance III spectrometers equipped with triple-resonance cryogenic TCI probes at 303 K. All the NMR data acquisition and processing parameters are outlined in Table S4. VimT protein samples were prepared by dialyzing 1 mL of isolated protein material obtained from anion exchange chromatography into 20 mM phosphate buffer (pH 6.5) for 2–3 h. Dialysis was continued with fresh buffer before overnight at room temperature. After dialysis, the protein concentration was determined by using absorbance measurements at 205 nm recorded with a 1 mm pathlength and a calculated extinction coefficient of 217,310 M^-1^ cm^-1^.^67^ The final protein NMR samples (44 μM) were prepared by adding neat D_2_O such that the final H_2_O:D_2_O was 90%:10% (v/v). All protein NMR spectra were processed using NMRPipe^68,69^ and further analyzed in SPARKY.^70^ Two-dimensional ^1^H–^15^N HSQC and three-dimensional HNCO, HNCACB, and HNCACOCB spectra of VimT were collected and analyzed to make sequence-specific assignments. The sequence-specific assignments of the VimT were determined manually following standard NMR chemical shift assignment strategies,^71,72^ and checked using the I-PINE web server^73^ (Table S5) through the POKY^74^ software platform. Tables S6 and S7 list the unambiguously assigned and unassigned peaks, respectively. The BMRB accession number for the VimT assignments is 52964.

^1^H–^15^N heteronuclear NOE spectra were recorded at 600 MHz ^1^H frequency using the standard Bruker pulse program, fhsqcf3gpph. Relaxation measurements (R_1_ and R_1ρ_ values) were recorded using heat compensated ^1^H–^15^N HSQC-based pulse program at 600 MHz ^1^H frequency.^75,76^ The R_1_ measurement used interleaved spectra recorded with relaxation delays of 40, 120, 200, 320, 520, 720, and 820 ms. The R_1ρ_ measurement used interleaved spectra recorded with relaxation delays of 1, 21, 41, 111, 131, and 151 ms and a B_1_ field of 1,400 Hz. ^15^N R_2_ rates were calculated from the R_1_ and R_1ρ_ values according to established methods.^47,48,76–78^ Peak intensities from the 2D spectra were obtained by removing overlapping signals in the ^1^H–^15^N HSQC spectra and integrating the remaining signals over +/-1 point in both dimensions using the NMRPipe software package.^68^ For the Cu(II) titration experiments, the protein sample (38 μM) was prepared following the same steps outlined above, and a 500 μM stock solution of CuCl_2_ was prepared in nanopure water. A range of 0 to 0.5 molar equivalents of Cu(II) were added directly into the same protein sample and allowed to incubate for 5–10 min at room temperature after each metal addition. A series of 2D ^1^H–^15^N HSQC were recorded, one for each Cu(II) equivalence (0, 0.1, 0.2, 0.3, 0.4, 0.5). The peaks in the spectra were filtered to remove overlapping signals and intensities obtained by integration with +/-2 point bounds in both spectral dimensions using NMRPipe.

## Abbreviations

The abbreviations used are

DIC: diisopropylcarbodiimide
DMF: dimethylformamide
DCM: dichloromethane
TFA: trifluoroacetic acid
TIPS: triisopropylsilane
IMAC: immobilized metal affinity chromatography, SDS-PAGE sodium dodecyl-sulfate polyacrylamide gel electrophoresis
TEV: tobacco etch virus
DTT: dithiothreitol
EDTA: ethylenediaminetetraacetic acid

## Data availability

Data presented in this study can be found in the manuscript or in the Supporting Information. Raw data files will be shared upon request.

## Supporting information

This article contains supporting information (38, 39, 40, 42, 50, 51, 68, 69, 70, 73).

## Acknowledgements

Dr. Steven L. McKnight generously provided the vimentin tail domain plasmid used in this work. We thank Dr. John Voss and Dr. Madhu Budamagunta for the training and use of the CD spectropolarimeter at the UC Davis Protein Structure and Dynamics Core Facility, and for their invaluable insights and project support. We also thank Dr. Upasana Sridharan and Dr. Blake Fonda for their assistance with NMR experiments and data processing, the UC Davis NMR and Campus Mass Spectrometry Facilities, and all the members of the Heffern and Murray laboratories for their helpful discussions.

## Author contributions

E.J.C investigation, data analysis, and writing–original draft; D.T.M and M.C.H conceptualization, supervision, funding acquisition, writing–review and editing.

## Funding and additional information

This work was supported by the National Institutes of Health and the UC Davis Chemical Biology Program training grant T32GM136597 to E.J.C., by the National Institute of General Medical Sciences of the National Institutes of Health under MIRA awards R35GM133684 to M.C.H and R35GM142892 to D.T.M, and by the National Science Foundation CAREER 2048265 to M.C.H. The UC Davis NMR Core Facility is supported in part by the National Science Foundation awards DBI-0722538, DBI-0079461, and DBI-0722538. The UC Davis Campus Mass Spectrometry Facility is partially funded by the National Institutes of Health Shared Instrumentation award S10OD025271. The content is solely the responsibility of the authors and does not necessarily represent the official views of the National Institutes of Health or the National Science Foundation.

