## Supplemental Figures and Tables for "Metal ion binding of vimentin tail domain fragments"

**Table of Contents**

Figure S1: Preparation of VimT

Figure S2: ESI-MS spectrum of nonacetylated VimTPep with Cu(II)

Figure S3: High-resolution ESI-MS spectra of VimTPep with Cu(II) and Zn(II)

Figure S4: High-resolution ESI-MS spectra of VimTPep with Ca(II) and Mg(II)

Table S1: Experimental values for calculating approx. binding affinity of Cu(II) to VimTPep

Table S2: Experimental values for calculating approx. binding affinity of Cu(II) to VimT

Figure S5: CD spectra of VimTPep with 0 to 10 equivalents of Cu(II)

Figure S6: 1D plots of CD spectra of VimTPep and VimT with Cu(II)

Table S3: Estimated secondary structure values of CD data obtained from BetStSel

Figure S7: CD spectra of VimT and Cu(II) at different protein concentrations

Figure S8: CD spectra of VimT with 0 to 0.5 equivalents of Cu(II) at pH 6.5

Table S4: NMR experimental data collection and processing parameters

Figure S9: AlphaFold 2.0 prediction of full-length vimentin monomer

Figure S10: NMR analysis of Cu(II) binding to VimT

Table S5: I-PINE automated assignment results of VimT

Table S6: Assigned VimT chemical shifts

Table S7: Unassigned VimT chemical shifts

Figure S11: AlphaFold 3.0 predicted models of vimentin with Cu(II) ions

References

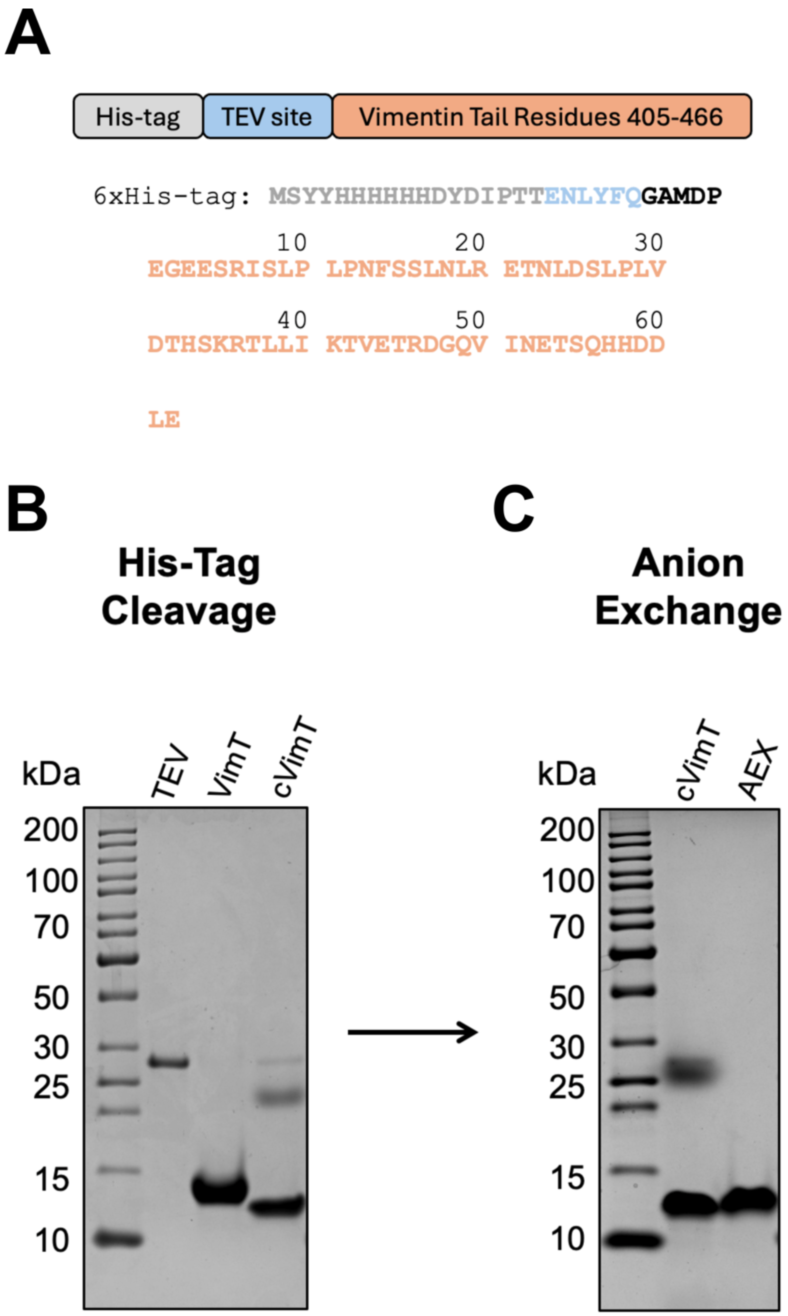


**Figure S1. Preparation of vimentin tail domain protein fragment.** (A) Amino acid sequence of VimT including the sequence of the purification 6xHis-tag. After the removal of the His-tag by TEV the following five non-native residues remain at the N-terminus: Gly, Ala, Met, Asp, Pro (GAMDP) for a total of 67 residues in the VimT fragment. SDS-PAGE gels of (B) vimentin tail domain protein fragment (VimT, ~14 kDa) before and after His-tag cleavage (cVimT, ~11 kDa) with TEV enzyme and (C) subsequent isolation of purified and His-tag cleaved VimT material by anion exchange chromatography (AEX). Upon TEV-cleavage, there is a presence of an additional cVimT protein band ~20 kDa that is removed in the AEX process. All protein experiments were carried out using the post-AEX VimT material.

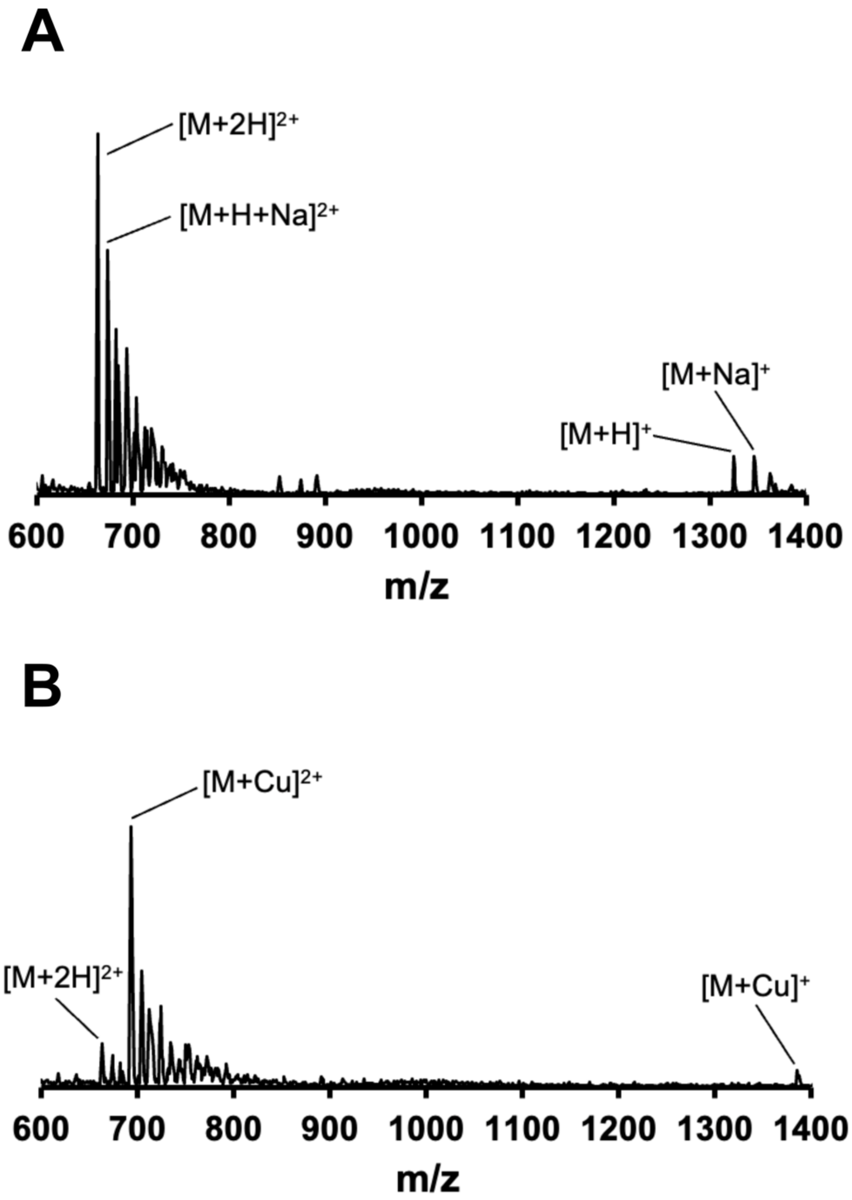


**Figure S2. ESI-MS spectra of non-acetylated VimTPep in the absence and presence of Cu(II).** Changes in the spectrum of (A) non-acetylated manually synthesized VimTPep (100 μM) are observed upon the addition of (B) 1 molar equivalent of Cu(II). Upon incubation with Cu(II), there is a large decrease in the signal intensity of mass peaks corresponding to VimTPep and the formation of VimTPep/Cu(II) adducts. The spectra support the direct binding interaction of VimTPep and Cu(II). Spectra were collected using our in-lab single quadrupole LC/MS instrument.

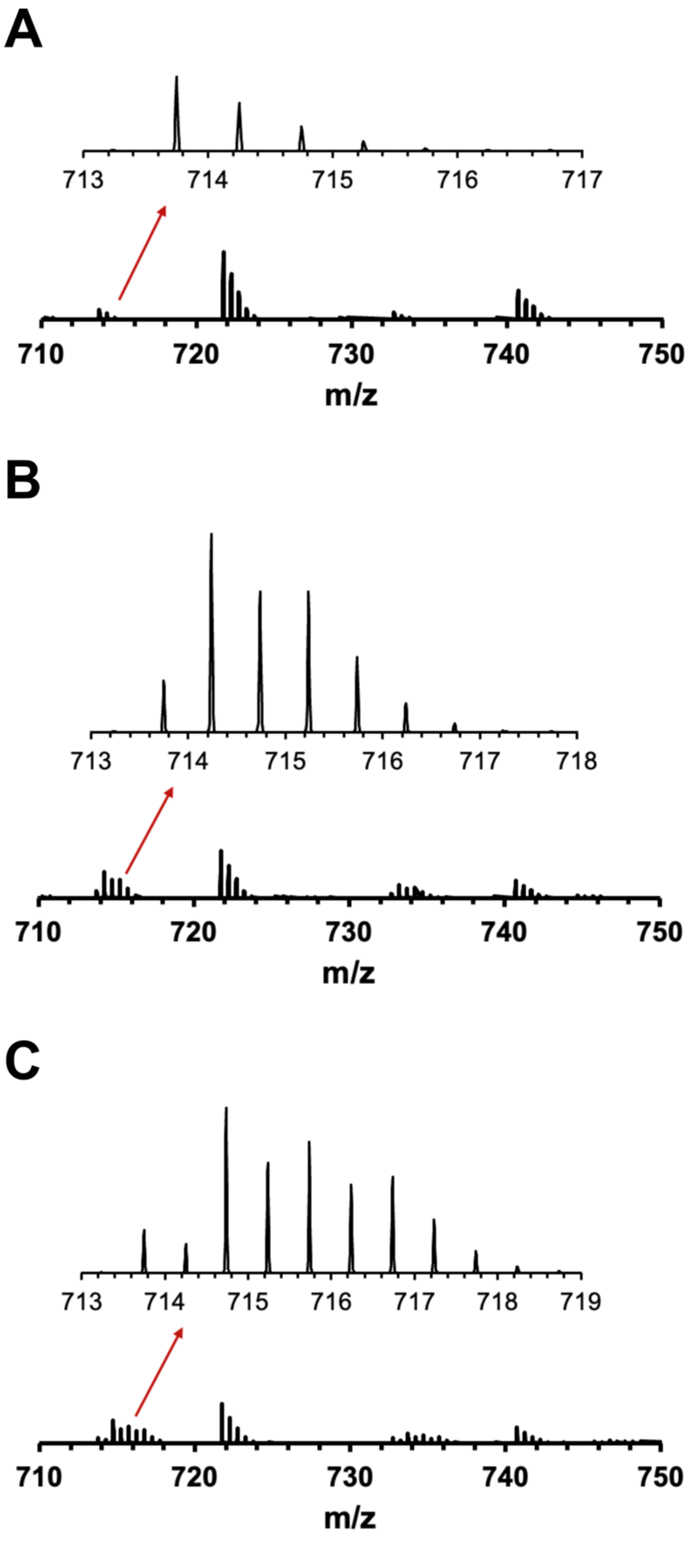


**Figure S3. High-resolution ESI-MS of VimTPep with Cu(II) and Zn(II).** The incubation of (A) VimTPep (50 μM) with 1 molar equivalent of (B) Cu(II) and (C) Zn(II) results in the formation of peptide-metal adducts and distinct isotopic mass distributions (red arrows), indicative of the direct metal interactions of Cu(II) and Zn(II) to VimTPep.

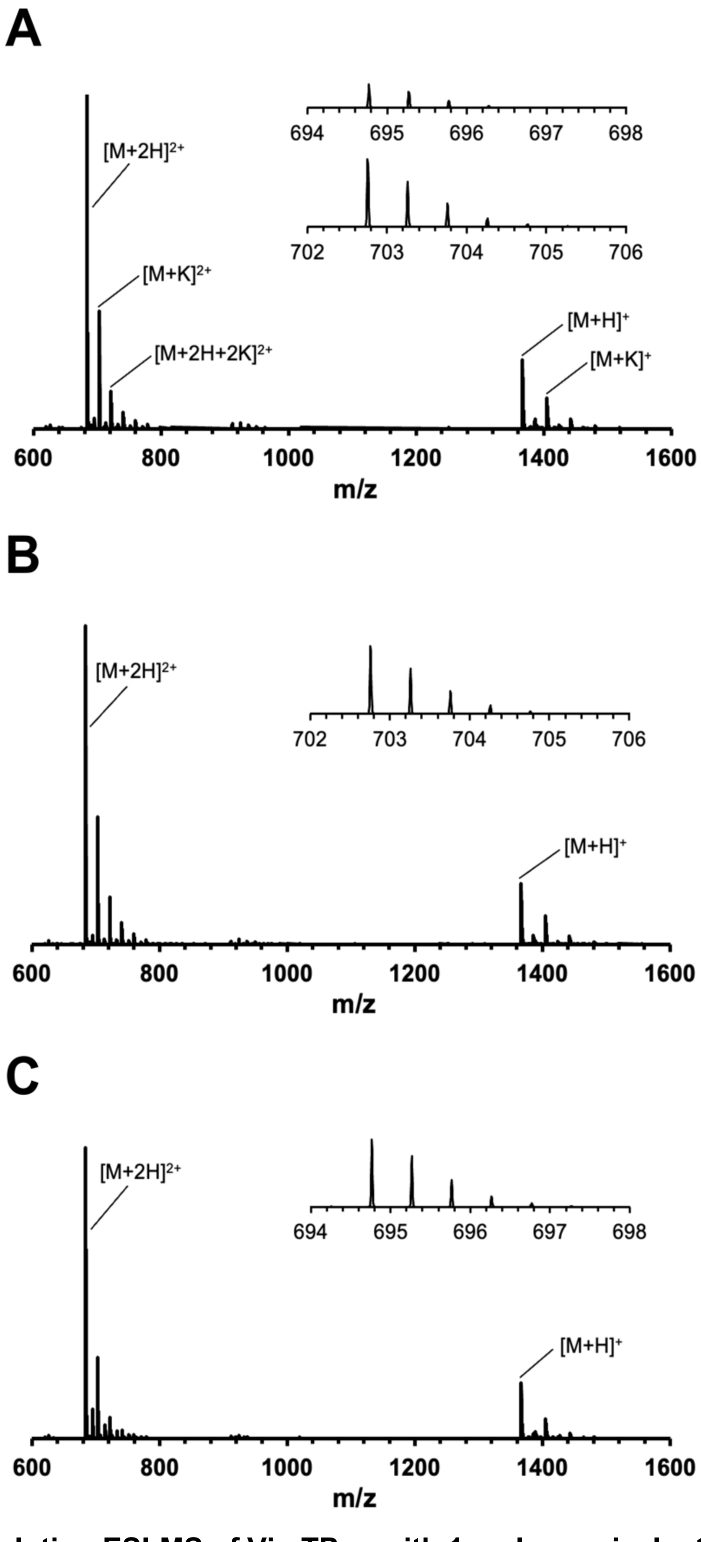


**Figure S4. High-resolution ESI-MS of VimTPep with 1 molar equivalent of Ca(II) and Mg(II).** The spectrum of (A) VimTPep (50 μM) with the addition of equimolar amounts of (B) Ca(II) and (C) Mg(II) largely resembles the spectrum of apo-peptide (no metal), with no apparent formation of peptide-metal adducts. (C) The incubation of VimTPep with Mg(II) intensifies the signals corresponding to mass peaks with *m/z* values between 694.5–697.5, but these masses align with those observed for apo-peptide.

**Table S1.** Experimental values and calculations for determining the formation constant of VimTPep/Cu(II) using the chromophoric ligand 1,10-phenanthroline (phen). The concentrations are in micromolar. Using the data from Figure 3A, the estimated binding affinity of Cu(II) to VimTPep is log *K* 7.5–8.7 (or $K_{d,app}$ = 2–32 nM). All calculations were performed following approaches previously reported.[1–3]

| [VimTPep] | $A_{265}$ | $[Cu(phen)_3]^{2+}$ | $[phen]_{free}$ | $[VimTPep]_{free}$ | [Cu(II)/VimTPep] | $K_{ex}$ ($x10^8$) | $K_d^{Cu(II)/VimTPep}$ | log $K_d$ Cu(II)/VimTPep |
|---|---|---|---|---|---|---|---|---|
| 0.0 | 0.8485 | 9.43 | 11.72 | 0.0 | 0.0 | ND | ND | ND |
| 9.1 | 0.8275 | 9.19 | 12.42 | 8.9 | 0.2 | 0.0355 | 2.81E-08 | 7.55 |
| 18.3 | 0.7395 | 8.22 | 15.35 | 17.1 | 1.2 | 0.1321 | 7.57E-09 | 8.12 |
| 27.7 | 0.6674 | 7.42 | 17.75 | 25.7 | 2.0 | 0.1877 | 5.33E-09 | 8.27 |
| 37.1 | 0.6045 | 6.72 | 19.85 | 34.4 | 2.7 | 0.2327 | 4.30E-09 | 8.37 |
| 46.7 | 0.5573 | 6.19 | 21.42 | 43.4 | 3.2 | 0.2577 | 3.88E-09 | 8.41 |
| 56.3 | 0.5109 | 5.68 | 22.97 | 52.5 | 3.8 | 0.2891 | 3.46E-09 | 8.46 |
| 65.9 | 0.4760 | 5.29 | 24.13 | 61.7 | 4.1 | 0.3058 | 3.27E-09 | 8.49 |
| 75.6 | 0.4433 | 4.93 | 25.22 | 71.1 | 4.5 | 0.3245 | 3.08E-09 | 8.51 |
| 85.3 | 0.4148 | 4.61 | 26.17 | 80.4 | 4.8 | 0.3403 | 2.94E-09 | 8.53 |
| 95.0 | 0.3831 | 4.26 | 27.23 | 89.8 | 5.2 | 0.3683 | 2.71E-09 | 8.57 |
| 104.8 | 0.3624 | 4.03 | 27.92 | 99.4 | 5.4 | 0.3769 | 2.65E-09 | 8.58 |
| 114.5 | 0.3398 | 3.78 | 28.67 | 108.9 | 5.7 | 0.3941 | 2.54E-09 | 8.60 |
| 124.3 | 0.3268 | 3.63 | 29.11 | 118.6 | 5.8 | 0.3919 | 2.55E-09 | 8.59 |
| 134.2 | 0.3069 | 3.41 | 29.77 | 128.1 | 6.0 | 0.4099 | 2.44E-09 | 8.61 |
| 144.0 | 0.2860 | 3.18 | 30.47 | 137.8 | 6.2 | 0.4350 | 2.30E-09 | 8.64 |
| 153.8 | 0.2726 | 3.03 | 30.91 | 147.4 | 6.4 | 0.4429 | 2.26E-09 | 8.65 |
| 163.7 | 0.2565 | 2.85 | 31.45 | 157.1 | 6.6 | 0.4620 | 2.16E-09 | 8.66 |
| 173.6 | 0.2472 | 2.75 | 31.76 | 166.9 | 6.7 | 0.4628 | 2.16E-09 | 8.67 |
| 183.4 | 0.2354 | 2.62 | 32.15 | 176.6 | 6.8 | 0.4742 | 2.11E-09 | 8.68 |
| 193.3 | 0.2271 | 2.52 | 32.43 | 186.4 | 6.9 | 0.4760 | 2.10E-09 | 8.68 |
| | | | | | | | Range | 7.5-8.7 |

**Table S2.** Experimental values and calculations for determining the formation constant of VimT/Cu(II) using the chromophoric ligand 1,10-phenanthroline (phen). The concentrations are in micromolar. From the data collected in Figure 3B, the estimated binding affinity of Cu(II) to VimT is log $K$ 7.5–8.6 (or $K_{d,app}$ = 2–27 nM). All calculations were performed following approaches previously reported.[1–3]

| [VimT] | $A_{265}$ | $[Cu(phen)_3]^{2+}$ | $[phen]_{free}$ | $[VimT]_{free}$ | [Cu(II)/VimT] | $K_{ex}$ (x10$^8$) | $K_d^{Cu(II)/VimT}$ | log $K_d^{Cu(II)/VimT}$ |
|---|---|---|---|---|---|---|---|---|
| 0.0 | 0.7534 | 8.37 | 14.89 | 0.0 | 0.0 | ND | ND | ND |
| 6.4 | 0.7421 | 8.25 | 15.26 | 6.2 | 0.1 | 0.0374 | 2.67E-08 | 7.57 |
| 12.8 | 0.6747 | 7.50 | 17.51 | 12.0 | 0.9 | 0.1706 | 5.86E-09 | 8.23 |
| 19.4 | 0.6054 | 6.73 | 19.82 | 17.7 | 1.6 | 0.2731 | 3.66E-09 | 8.44 |
| 26.0 | 0.5559 | 6.18 | 21.47 | 23.8 | 2.2 | 0.3204 | 3.12E-09 | 8.51 |
| 32.7 | 0.5214 | 5.79 | 22.62 | 30.1 | 2.6 | 0.3345 | 2.99E-09 | 8.52 |
| 39.4 | 0.4926 | 5.47 | 23.58 | 36.5 | 2.9 | 0.3421 | 2.92E-09 | 8.53 |
| 46.1 | 0.4617 | 5.13 | 24.61 | 42.9 | 3.2 | 0.3627 | 2.76E-09 | 8.56 |
| 52.9 | 0.4424 | 4.92 | 25.25 | 49.4 | 3.5 | 0.3592 | 2.78E-09 | 8.56 |
| 59.7 | 0.4211 | 4.68 | 25.96 | 56.0 | 3.7 | 0.3660 | 2.73E-09 | 8.56 |
| 66.5 | 0.4006 | 4.45 | 26.65 | 62.6 | 3.9 | 0.3749 | 2.67E-09 | 8.57 |
| 73.3 | 0.3849 | 4.28 | 27.17 | 69.2 | 4.1 | 0.3756 | 2.66E-09 | 8.57 |
| 80.2 | 0.3708 | 4.12 | 27.64 | 75.9 | 4.3 | 0.3757 | 2.66E-09 | 8.57 |
| 87.0 | 0.3539 | 3.93 | 28.20 | 82.6 | 4.4 | 0.3855 | 2.59E-09 | 8.59 |
| 93.9 | 0.3463 | 3.85 | 28.46 | 89.4 | 4.5 | 0.3742 | 2.67E-09 | 8.57 |
| 100.8 | 0.3372 | 3.75 | 28.76 | 96.2 | 4.6 | 0.3692 | 2.71E-09 | 8.57 |
| | | | | | | | Range | 7.5-8.6 |

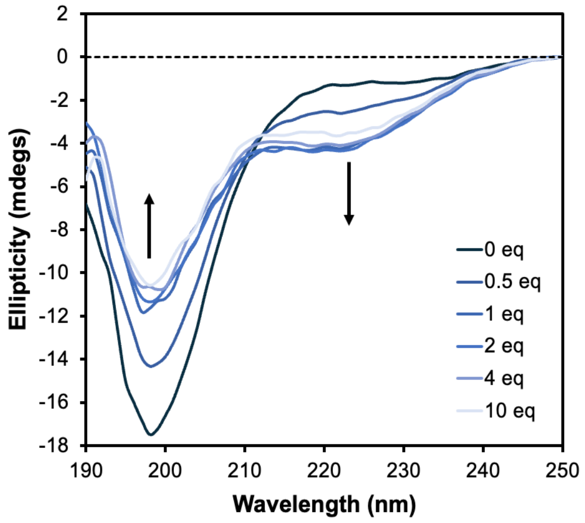


**Figure S5. Circular dichroism spectra of VimTPep (100 μM) in 20 mM phosphate buffer (pH 7.4).** Changes in the random coil confirmation of peptide are observed upon the addition of 0 to 10 molar equivalents (eq) of Cu(II), demonstrated by the reduction of ellipticity at 198 nm and increase of ellipticity between 220 and 230 nm. Reduction in the negative ellipticity of the peptide is greatest between 0 to 1 eq Cu(II), with minimal changes in spectra between 1 to 10 eq Cu(II).

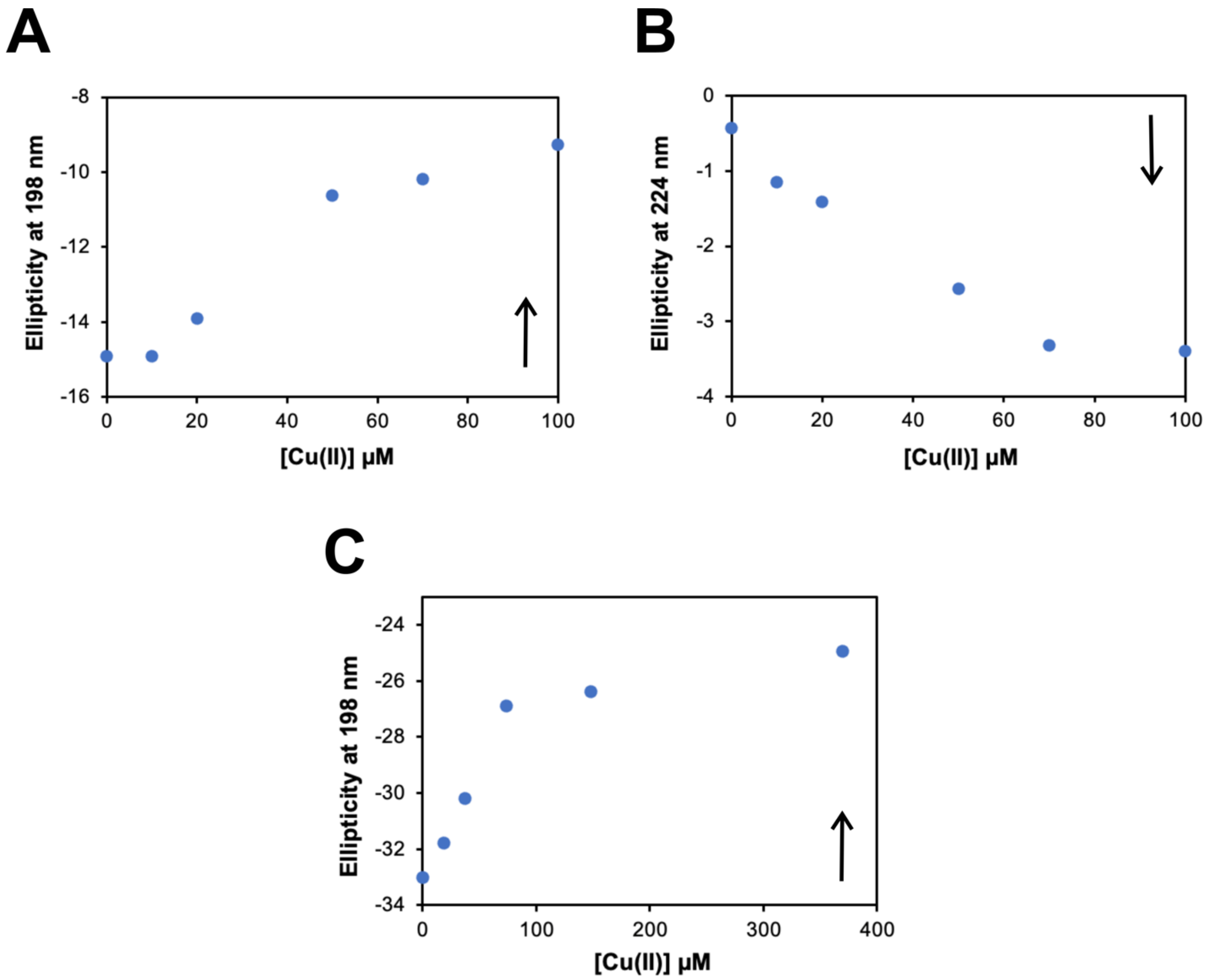


**Figure S6. Circular dichroism one-dimensional plots of VimTPep and VimT spectra with Cu(II).** Plots of the ellipticity values of VimTPep (100 µM) at (A) 198 nm and (B) 224 nm as a function of Cu(II) concentration, corresponding to 0–1 Cu(II) equivalent range. (C) A plot of VimT (~37 µM) ellipticity values at 198 nm as a function of Cu(II) concentrations, showing the minimal decrease in the negative ellipticity of VimT at 0–10 Cu(II) equivalents.

**Table S3.** Summary of the estimated secondary structure content of VimTPep and VimT with increasing addition of Cu(II) obtained from fitting the CD spectra of both tail domain fragments by using the BetStSel[4] algorithm. The "Others" structural content includes disordered segments, bend, loop, $\pi$-helix, $3_{10}$ helix, and $\beta$-bridge regions. The web server is available at https://bestsel.elte.hu.

| Estimated Secondary Structure Content (%) | | | | | | | | | | | | |
|---|---|---|---|---|---|---|---|---|---|---|---|---|
| | **VimTPep** | | | | | | **VimT** | | | | | |
| Cu(II) Equivalent (eq) | 0 eq | 0.1 eq | 0.2 eq | 0.5 eq | 0.7 eq | 1 eq | 0 eq | 0.5 eq | 1.0 eq | 2 eq | 4 eq | 10 eq |
| Helix | 0.0 | 0.0 | 0.0 | 0.0 | 0.3 | 1 | 0.1 | 0.2 | 0.9 | 1.1 | 1.8 | 1.9 |
| Antiparallel | 34.3 | 32.8 | 32.6 | 32.6 | 31 | 30.8 | 30.4 | 30.4 | 30.6 | 30.1 | 30.0 | 29.9 |
| Parallel | 0.0 | 0.0 | 0.0 | 0.0 | 0.0 | 0.0 | 0.0 | 0.0 | 0.0 | 0.0 | 0.0 | 0.0 |
| Turn | 18.5 | 18.0 | 17.7 | 16.8 | 16.5 | 16.8 | 17.8 | 17.8 | 17.4 | 17.0 | 16.7 | 16.6 |
| Others | 47.2 | 49.2 | 49.6 | 50.6 | 52.2 | 51.5 | 51.6 | 51.7 | 51.1 | 51.8 | 51.5 | 51.5 |

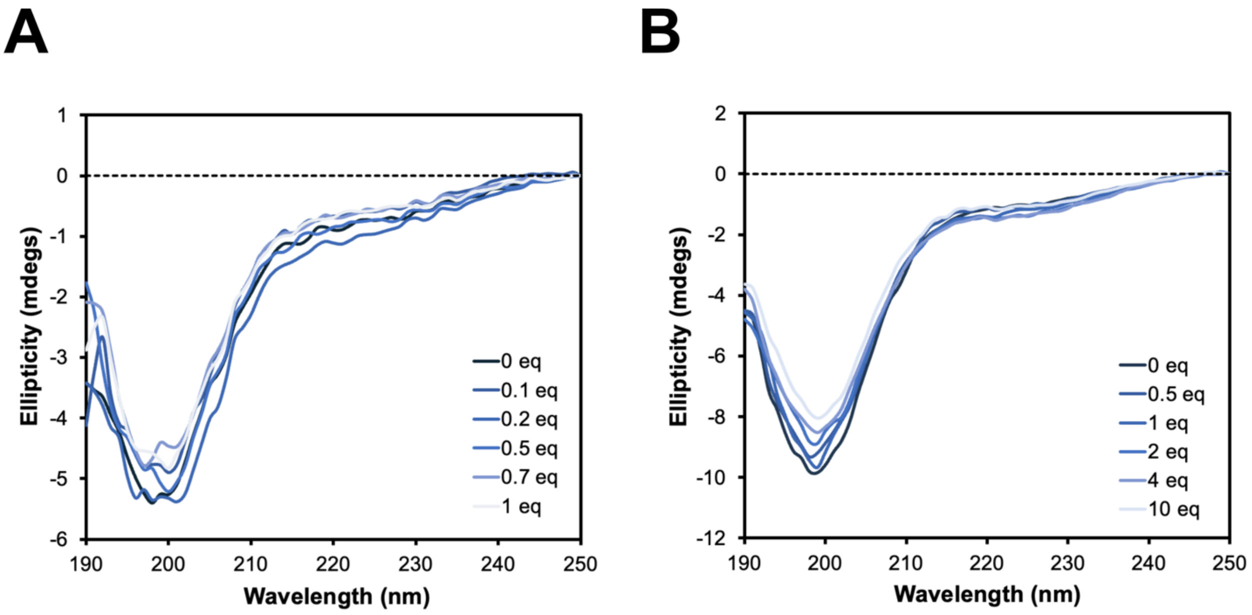


**Figure S7. Circular dichroism (CD) spectra of VimT with Cu(II) at varying protein concentrations.** Increasing concentration of VimT from (A) 6 μM (or 0.04 mg/mL) to (B) 14.3 μM (or 0.107 mg/mL) resulted in better resolved spectral features and improved the overall CD signal, with the optimal the concentration of VimT for CD determined to be ~37 μM (or ~0.3 mg/mL).

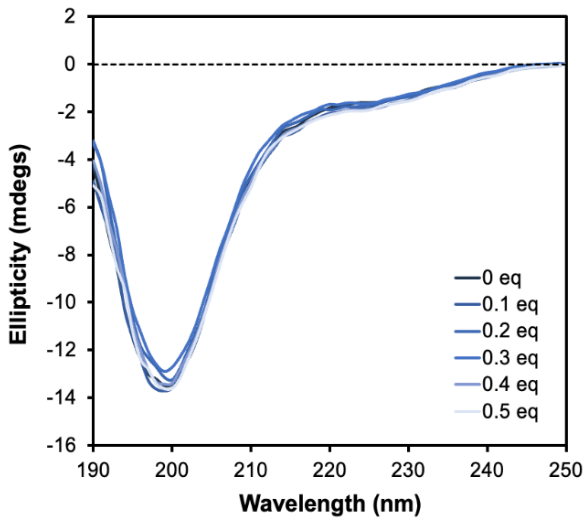


**Figure S8. Circular dichroism spectra of VimT protein (14.4 μM) with the addition of 0 to 0.5 molar equivalents (eq) of Cu(II) at pH 6.5.** Increasing amounts of Cu(II) leads to minimal decrease in the negative ellipticity at 198 nm and minimal changes overall, indicating that the protein fragment remains in a random coil conformation.

**Table S4.** Solution NMR vimentin tail domain protein data collection and processing parameters.

| Spectrum | Acquisition Parameters[a] | Processing Parameters[b] |
|---|---|---|
| 2D $^{1}H$–$^{15}N$ HSQC (for assignments) | 600 MHz Bruker AVANCE III<br>ns = 32; $\tau_{aq}$ = 159.7 ms with 3072 points; $\tau_{t1}$ = 90.2 ms with 512 States-TPPI points; $\nu_{1H\text{-}carr}$ = 4.694 ppm; $\nu_{15N\text{-}carr}$ = 117.000 ppm; NusAMOUNT = 50% random point sampling; Temp = 298.2 K | NMRPipe IST processing with ist2D.com and standard NUS parameters |
| 2D $^{1}H$–$^{15}N$ HSQC (for Cu titrations) | 800 MHz Bruker AVANCE III<br>ns = 16; $\tau_{aq}$ = 79.9 ms with 2048 points; $\tau_{t1}$ = 45.1 ms with 256 States-TPPI points; $\nu_{1H\text{-}carr}$ = 4.697 ppm; $\nu_{15N\text{-}carr}$ = 117.000 ppm;<br>Temp = 295.8 K | Reference correction in Sparky:<br>$^{1}H$: 0.07 ppm (for 0.1 equivalent Cu spectrum only) |
| 3D HNCO | 600 MHz Bruker AVANCE III<br>ns = 8; $\tau_{aq}$ = 170.4 ms with 3072 points; $\tau_{t1\,N}$ = 15.4 ms with 60 States-TPPI points; $\tau_{t2\,C}$ = 21.2 ms with 128 States-TPPI points; $\nu_{1H\text{-}carr}$ = 4.700 ppm; $\nu_{15N\text{-}carr}$ = 118.010 ppm; $\nu_{13C\text{-}carr}$ = 174.015 ppm; NusAMOUNT = 40% random point sampling; Temp = 298.2 K | NMRPipe IST processing with ist3D.com and standard NUS parameters<br><br>Reference correction in Sparky: $^{1}H$: 0.01ppm |
| 3D HNCACB | 600 MHz Bruker AVANCE III<br>ns = 8; $\tau_{aq}$ = 170.4 ms with 3072 points; $\tau_{t1\,N}$ = 21.6 ms with 84 States-TPPI points; $\tau_{t2\,C}$ = 7.2 ms with 152 States-TPPI points; $\nu_{1H\text{-}carr}$ = 4.700 ppm; $\nu_{15N\text{-}carr}$ = 118.010 ppm; $\nu_{13C\text{-}carr}$ = 39.003 ppm; NusAMOUNT = 38% random point sampling; Temp = 298.2 K | NMRPipe IST processing with ist3D.com and standard NUS parameters |
| 3D CBCA(CO)NH | 600 MHz Bruker AVANCE III<br>ns = 8; $\tau_{aq}$ = 170.4 ms with 3072 points; $\tau_{t1\,N}$ = 15.4 ms with 60 States-TPPI points; $\tau_{t2\,C}$ = 5.7 ms with 120 States-TPPI points; $\nu_{1H\text{-}carr}$ = 4.693 ppm; $\nu_{15N\text{-}carr}$ = 118.010 ppm; $\nu_{13C\text{-}carr}$ = 39.003 ppm; NusAMOUNT = 45% of random point sampling; Temp = 298.2 K | NMRPipe IST processing with ist3D.com and standard NUS parameters<br><br>Reference correction in Sparky:<br>$^{13}C$: 1.03 ppm |

[a] ns, number of scans; $\tau_{aq}$, total acquisition time for the direct dimension; $\tau_{tX}$, total acquisition time for the indirect dimension x; $\nu_{X\text{-}carr}$, center of the frequency range for nucleus x. The proton pulse length was calibrated and optimized by running a 1D proton experiment and using the Bruker "pulsecal" command.

[b]All spectra were processed with NMRPipe and using standard NUS parameters where applicable.[5,6] Reference corrections were applied to align spectra to the peaks found in the HSQC used for assignments or for Cu titrations (0 Cu equivalent spectrum).

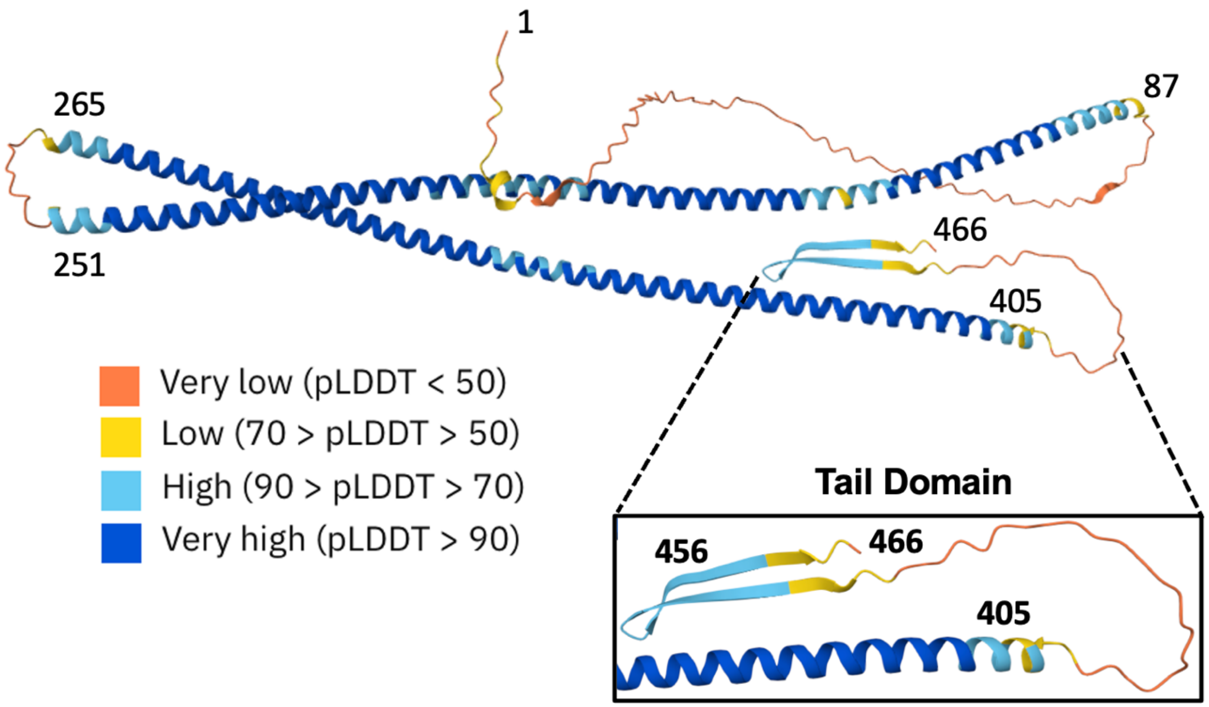


**Figure S9. AlphaFold 2.0[7] prediction of full-length vimentin structure (P08670)**. The inset view shows the tail domain region investigated in this work spanning residues 405–466, with primarily low (yellow) to very low (orange) assigned per-residue model confidence scores (pLDDT). A beta site containing the conserved beta-turn (449–452) is predicted with reasonably high confidence for residues 443–460.

**Table S5.** I-PINE[8] automated assignments of vimentin tail domain protein NMR resonances. Column 1 is the residue number for the 67 residues of VimT, with the native residues beginning at residue number 405 (GLU) and negative numbers representing non-native residues from the purification tag. The I-PINE results provided a total of four probabilistic assignments (P(H,N)) based on the submitted NMR chemical shifts and VimT protein sequence. The probability values range from 0 (low) to 1 (high). The values corresponding to the two highest assignment probabilities, columns 3 and 9, are shown for clarity. The assignment with the highest probability lists the $^{1}$H, $^{15}$N, $^{13}$CO, $^{13}$C$\alpha$ and $^{13}$C$\beta$ chemical shifts (columns 4–8), with subsequent probabilities reporting only the $^{1}$H and $^{15}$N chemical shifts. The final column indicates the probability of each residue having no assignment (P(no assignment)). Missing values are displayed as "999". The I-PINE web server can be accessed online at http://i-pine.nmrfam.wisc.edu.

| Residue Number | Residue Name | P (H,N) | H | N | CO | CA | CB | P (H,N) | H | N | P (no assignment) |
|---|---|---|---|---|---|---|---|---|---|---|---|
| –5 | GLY | 0 | 0 | 0 | 176.7 | 999 | 0 | 0 | 0 | 0 | 0 |
| –4 | ALA | 0.045 | 8.252 | 121.81 | 177.9 | 53.1 | 19.3 | 0 | 0 | 0 | 0.954 |
| –3 | MET | 0.805 | 8.489 | 117.8 | 0 | 0 | 0 | 0.002 | 8.252 | 121.81 | 0.193 |
| –2 | ASP | 0.166 | 8.441 | 110 | 0 | 999 | 999 | 0.032 | 8.252 | 121.81 | 0.785 |
| –1 | PRO | 0 | 0 | 0 | 177.27 | 63.5 | 32 | 0 | 0 | 0 | 0 |
| 405 | GLU | 1 | 8.535 | 120.17 | 177.17 | 56.7 | 30.1 | 0 | 0 | 0 | 0 |
| 406 | GLY | 1 | 8.289 | 109.52 | 174.43 | 45.3 | 0 | 0 | 0 | 0 | 0 |
| 407 | GLU | 1 | 8.336 | 120.89 | 177.09 | 56.9 | 30.4 | 0 | 0 | 0 | 0 |
| 408 | GLU | 1 | 8.641 | 121.42 | 176.88 | 57.2 | 29.9 | 0 | 0 | 0 | 0 |
| 409 | SER | 1 | 8.251 | 116.22 | 174.68 | 58.7 | 63.5 | 0 | 0 | 0 | 0 |
| 410 | ARG | 1 | 8.19 | 122.87 | 176.21 | 56.1 | 30.7 | 0 | 0 | 0 | 0 |
| 411 | ILE | 1 | 8.027 | 121.14 | 176.1 | 61 | 38.6 | 0 | 0 | 0 | 0 |
| 412 | SER | 1 | 8.319 | 120.04 | 173.83 | 57.8 | 63.7 | 0 | 0 | 0 | 0 |
| 413 | LEU | 1 | 8.196 | 125.86 | 0 | 999 | 41.8 | 0 | 0 | 0 | 0 |
| 414 | PRO | 0 | 0 | 0 | 176.53 | 62.6 | 31.8 | 0 | 0 | 0 | 0 |
| 415 | LEU | 0.63 | 8.267 | 123.67 | 0 | 999 | 41.8 | 0.006 | 8.252 | 121.81 | 0.363 |
| 416 | PRO | 0 | 0 | 0 | 176.53 | 999 | 999 | 0 | 0 | 0 | 0 |
| 417 | ASN | 0.235 | 8.267 | 123.67 | 0 | 999 | 999 | 0.051 | 8.031 | 121.5 | 0.694 |
| 418 | PHE | 0.366 | 8.441 | 110 | 176.18 | 999 | 999 | 0.099 | 8.355 | 118.13 | 0.457 |
| 419 | SER | 0.155 | 8.031 | 121.5 | 175.46 | 999 | 999 | 0.043 | 8.252 | 121.81 | 0.777 |
| 420 | SER | 0.157 | 7.989 | 121.33 | 174.58 | 58.6 | 63.5 | 0.035 | 7.991 | 115.29 | 0.787 |
| 421 | LEU | 1 | 8.078 | 123.18 | 177.03 | 55.5 | 42.3 | 0 | 0 | 0 | 0 |
| 422 | ASN | 1 | 8.314 | 118.71 | 175.69 | 53.2 | 38.5 | 0 | 0 | 0 | 0 |
| 423 | LEU | 1 | 8.113 | 122.41 | 177.31 | 55.3 | 42.1 | 0 | 0 | 0 | 0 |
| 424 | ARG | 1 | 8.229 | 120.95 | 176.29 | 56.1 | 30.7 | 0 | 0 | 0 | 0 |
| 425 | GLU | 1 | 8.392 | 121.53 | 176.57 | 56.7 | 30.2 | 0 | 0 | 0 | 0 |
| 426 | THR | 1 | 8.129 | 114.59 | 174.1 | 61.6 | 69.8 | 0 | 0 | 0 | 0 |
| 427 | ASN | 1 | 8.499 | 121.17 | 175.27 | 53.2 | 38.7 | 0 | 0 | 0 | 0 |
| 428 | LEU | 1 | 8.293 | 122.69 | 177.36 | 55.5 | 42.1 | 0 | 0 | 0 | 0 |
| 429 | ASP | 1 | 8.275 | 120.05 | 176.18 | 54.6 | 40.9 | 0 | 0 | 0 | 0 |
| 430 | SER | 1 | 8.011 | 114.9 | 174.07 | 57.9 | 63.8 | 0 | 0 | 0 | 0 |
| 431 | LEU | 1 | 8.092 | 125 | 0 | 999 | 41.8 | 0 | 0 | 0 | 0 |
| 432 | PRO | 0 | 0 | 0 | 0 | 53.2 | 999 | 0 | 0 | 0 | 0 |
| 433 | LEU | 1 | 8.113 | 122.41 | 177.44 | 55.2 | 42.2 | 0 | 0 | 0 | 0 |
| 434 | VAL | 1 | 8.054 | 120.69 | 175.64 | 62 | 32.9 | 0 | 0 | 0 | 0 |
| 435 | ASP | 1 | 8.412 | 124.18 | 175.69 | 54.4 | 41.2 | 0 | 0 | 0 | 0 |
| 436 | THR | 1 | 8.396 | 120.34 | 176.05 | 999 | 999 | 0 | 0 | 0 | 0 |
| 437 | HIS | 0.948 | 7.991 | 115.29 | 176.63 | 56.7 | 30.2 | 0.001 | 8.252 | 121.81 | 0.05 |
| 438 | SER | 0.974 | 8.151 | 114.83 | 0 | 55.6 | 999 | 0.009 | 7.989 | 121.33 | 0.011 |
| 439 | LYS | 0.977 | 8.497 | 122.06 | 176.72 | 999 | 999 | 0.002 | 7.989 | 121.33 | 0.019 |
| 440 | ARG | 0.983 | 8.256 | 121.64 | 176.49 | 56.4 | 30.8 | 0.017 | 8.252 | 121.81 | 0 |

| | | | | | | | | | | | |
|---|---|---|---|---|---|---|---|---|---|---|---|
| 441 | THR | 1 | 8.156 | 115.85 | 174.21 | 62.1 | 69.6 | 0 | 0 | 0 | 0 |
| 442 | LEU | 1 | 8.193 | 124.87 | 176.75 | 55.2 | 42.4 | 0 | 0 | 0 | 0 |
| 443 | LEU | 1 | 8.173 | 123.5 | 176.79 | 55 | 42.2 | 0 | 0 | 0 | 0 |
| 444 | ILE | 1 | 8.053 | 122.44 | 175.91 | 60.7 | 38.5 | 0 | 0 | 0 | 0 |
| 445 | LYS | 1 | 8.402 | 125.98 | 176.38 | 55.9 | 33.1 | 0 | 0 | 0 | 0 |
| 446 | THR | 1 | 8.226 | 116.85 | 174.31 | 61.7 | 69.8 | 0 | 0 | 0 | 0 |
| 447 | VAL | 1 | 8.236 | 122.32 | 175.78 | 62.1 | 33 | 0 | 0 | 0 | 0 |
| 448 | GLU | 1 | 8.539 | 124.97 | 176.72 | 56.3 | 30.4 | 0 | 0 | 0 | 0 |
| 449 | THR | 1 | 8.264 | 115.74 | 176.7 | 61.6 | 69.7 | 0 | 0 | 0 | 0 |
| 450 | ARG | 0.585 | 8.252 | 121.81 | 176.05 | 56 | 30.9 | 0.007 | 8.256 | 121.64 | 0.399 |
| 451 | ASP | 1 | 8.463 | 121.63 | 176.67 | 54.5 | 40.9 | 0 | 0 | 0 | 0 |
| 452 | GLY | 1 | 8.375 | 108.73 | 174.1 | 45.5 | 0 | 0 | 0 | 0 | 0 |
| 453 | GLN | 1 | 8.085 | 119.73 | 175.46 | 999 | 999 | 0 | 0 | 0 | 0 |
| 454 | VAL | 0.358 | 7.989 | 121.33 | 176.05 | 62.4 | 32.6 | 0.087 | 8.031 | 121.5 | 0.545 |
| 455 | ILE | 0.998 | 8.337 | 125.41 | 175.63 | 60.7 | 999 | 0 | 0 | 0 | 0.001 |
| 456 | ASN | 1 | 8.53 | 123.22 | 175.06 | 53 | 39 | 0 | 0 | 0 | 0 |
| 457 | GLU | 1 | 8.538 | 122.18 | 176.71 | 56.9 | 30.3 | 0 | 0 | 0 | 0 |
| 458 | THR | 1 | 8.27 | 114.69 | 174.71 | 62.1 | 69.7 | 0 | 0 | 0 | 0 |
| 459 | SER | 1 | 8.295 | 118 | 174.33 | 58.4 | 63.7 | 0 | 0 | 0 | 0 |
| 460 | GLN | 1 | 8.331 | 121.82 | 175.77 | 55.4 | 29.6 | 0 | 0 | 0 | 0 |
| 461 | HIS | 0.996 | 8.287 | 122.41 | 175.13 | 55.3 | 38.6 | 0.002 | 7.989 | 121.33 | 0 |
| 462 | HIS | 0.995 | 8.192 | 120.98 | 174.26 | 55.6 | 30.2 | 0 | 0 | 0 | 0.004 |
| 463 | ASP | 1 | 8.497 | 122.06 | 175.65 | 54.4 | 41.2 | 0 | 0 | 0 | 0 |
| 464 | ASP | 1 | 8.387 | 120.58 | 175.98 | 54.4 | 40.9 | 0 | 0 | 0 | 0 |
| 465 | LEU | 1 | 8.18 | 122.01 | 176.54 | 55.2 | 42.3 | 0 | 0 | 0 | 0 |
| 466 | GLU | 1 | 7.863 | 126.25 | 0 | 0 | 0 | 0 | 0 | 0 | 0 |

**Table S6.** Assigned vimentin tail domain protein NMR chemical shifts. The first column corresponds to the residue number with respect to the full-length vimentin protein, and S/N represents the signal-to-noise for each signal in the $^{1}H$–$^{15}N$ HSQC spectrum based on the peak intensities calculated by NMRFAM-SPARKY.[9] The NMR chemical shift assignments were determined from the values obtained from a combination of HSQC, HNCACB, HNCO, and HN(CO)CACB experiments. In the cases where the same signal was observed in different spectra (e.g. N sites in the HSQC and HNCACB), the values obtained from the HSQC spectrum are reported. Negative numbers represent non-native residues from the purification tag.

| Residue Number | Residue | CA | CB | CO | N | HN | S/N |
|---|---|---|---|---|---|---|---|
| –5 | G | 45.246 | | 177.093 | 109.999 | 8.441 | 24 |
| –4 | A | | | | | | |
| –3 | M | | | 177.900 | 117.798 | 8.489 | 59 |
| –2 | D | | | | 122.41 | 8.113 | 344 |
| –1 | P | 63.562 | 32.073 | | | | |
| 405 | E | 56.784 | 30.092 | 177.271 | 120.172 | 8.535 | 424 |
| 406 | G | 45.338 | | 177.174 | 109.517 | 8.289 | 231 |
| 407 | E | 56.982 | 30.359 | 174.428 | 120.895 | 8.336 | 242 |
| 408 | E | 57.281 | 30.011 | 177.081 | 121.419 | 8.641 | 264 |
| 409 | S | 58.694 | 63.537 | 176.884 | 116.219 | 8.251 | 153 |
| 410 | R | 56.173 | 30.707 | 174.683 | 122.870 | 8.190 | 112 |
| 411 | I | 61.251 | 38.800 | 176.213 | 121.143 | 8.027 | 195 |
| 412 | S | 58.056 | 63.743 | 176.098 | 120.044 | 8.319 | 107 |
| 413 | L | 52.938 | 41.944 | 173.817 | 125.865 | 8.196 | 219 |
| 414 | P | 62.687 | 31.862 | | | | |
| 415 | L | 52.924 | 41.713 | 176.547 | 123.675 | 8.267 | 349 |
| 416 | P | | | | | | |
| 417 | N | | | | | | |
| 418 | F | | | | | | |
| 419 | S | | | | | | |
| 420 | S | | | | | | |
| 421 | L | 55.687 | 42.337 | 174.584 | 123.179 | 8.078 | 111 |
| 422 | N | 53.253 | 38.527 | 177.037 | 118.714 | 8.314 | 84 |
| 423 | L | 55.362 | 42.162 | 175.118 | 120.978 | 8.192 | 46 |
| 424 | R | 56.218 | 30.765 | 177.310 | 120.948 | 8.229 | 156 |
| 425 | E | 56.591 | 30.286 | 176.306 | 121.532 | 8.392 | 150 |
| 426 | T | 61.894 | 69.993 | 176.557 | 114.592 | 8.129 | 128 |
| 427 | N | 53.280 | 38.683 | 174.143 | 121.175 | 8.499 | 61 |
| 428 | L | 55.478 | 42.167 | 175.257 | 122.693 | 8.293 | 207 |
| 429 | D | 54.643 | 41.067 | 177.357 | 120.052 | 8.275 | 226 |
| 430 | S | 58.092 | 63.806 | 176.183 | 114.902 | 8.011 | 155 |
| 431 | L | 53.318 | 41.616 | 174.074 | 125.004 | 8.092 | 302 |

| | | | | | | | |
|---|---|---|---|---|---|---|---|
| 432 | P | | | | | | |
| 433 | L | 55.400 | 42.100 | 175.049 | 122.4 | 8.114 | |
| 434 | V | 62.071 | 33.027 | 177.433 | 120.685 | 8.054 | 269 |
| 435 | D | 54.103 | 41.365 | 175.622 | 124.180 | 8.412 | 123 |
| 436 | T | | | 175.652 | 120.57 | 8.386 | 13 |
| 437 | H | | | | | | |
| 438 | S | | | | | | |
| 439 | K | | | | | | |
| 440 | R | | | | | | |
| 441 | T | | 69.649 | 176.493 | 115.851 | 8.156 | 39 |
| 442 | L | 55.167 | 42.319 | 174.222 | 124.866 | 8.193 | 61 |
| 443 | L | 55.167 | 42.319 | 176.748 | 123.5 | 8.173 | 102 |
| 444 | I | 60.836 | 38.733 | 176.794 | 122.436 | 8.053 | 254 |
| 445 | K | 56.233 | 33.248 | 175.902 | 125.982 | 8.402 | 117 |
| 446 | T | 61.842 | 69.924 | 176.379 | 116.848 | 8.226 | 104 |
| 447 | V | 62.073 | 32.973 | 174.313 | 122.317 | 8.236 | 218 |
| 448 | E | 56.297 | 30.662 | 175.768 | 124.966 | 8.539 | 217 |
| 449 | T | 61.656 | 70.164 | 176.437 | 115.741 | 8.264 | 127 |
| 450 | R | | | | | | |
| 451 | D | 54.631 | 41.018 | 176.037 | 121.630 | 8.463 | 105 |
| 452 | G | 45.504 | | 176.667 | 108.726 | 8.375 | 125 |
| 453 | Q | 55.652 | 29.818 | 174.156 | 119.726 | 8.085 | 206 |
| 454 | V | 62.734 | 32.490 | | 122.410 | 8.287 | |
| 455 | I | 60.835 | 38.785 | 176.056 | 125.408 | 8.337 | 357 |
| 456 | N | 53.102 | 39.091 | 175.646 | 123.222 | 8.530 | 151 |
| 457 | E | 56.954 | 30.350 | 175.044 | 122.18 | 8.538 | 138 |
| 458 | T | 62.087 | 69.799 | 176.713 | 114.694 | 8.270 | 100 |
| 459 | S | 58.496 | 63.736 | 174.701 | 117.996 | 8.295 | 53 |
| 460 | Q | 55.886 | 29.612 | 174.326 | 121.816 | 8.331 | 59 |
| 461 | H | | | | | | |
| 462 | H | | | | | | |
| 463 | D | 54.441 | 41.320 | 174.283 | 122.06 | 8.497 | 71 |
| 464 | D | 54.500 | 41.121 | 175.698 | 120.345 | 8.396 | 344 |
| 465 | L | 55.078 | 42.446 | 175.983 | 122.009 | 8.180 | 465 |
| 466 | E | 58.034 | 31.247 | 176.544 | 126.250 | 7.863 | 595 |

**Table S6.** Unassigned vimentin tail domain protein NMR chemical shifts. The values for the unassigned signals, shown in gray in Figure 6, were obtained from the 2D and 3D NMR experiments including the chemical shift values for the "*i-1*" residues. S/N represents the signal-to-noise values for the peaks in the $^{1}H–^{15}N$ HSQC spectrum analyzed by NMRFAM-SPARKY.[9]

| **N** | **CA** | **CB** | **$H_N$** | **CA (i-1)** | **CB (i-1)** | **CO (i-1)** | **S/N** |
|---|---|---|---|---|---|---|---|
| 114.829 | 69.50 | | 8.151 | | 41.27 | 176.634 | 36 |
| 115.286 | | | 7.991 | | | 176.047 | 26 |
| 118.127 | | 38.68 | 8.355 | 63.07 | 31.70 | 176.323 | 53 |
| 121.316 | | 42.20 | 7.933 | | | 174.668 | 33 |
| 121.335 | | | 7.989 | | | 175.463 | 25 |
| 121.504 | | | 8.031 | | | 176.175 | 42 |
| 121.636 | | 56.43 | 8.256 | | | 176.723 | 35 |
| 121.812 | 56.43 | 41.35 | 8.252 | | | 176.697 | 46 |
| 121.989 | | | 8.693 | | | 176.353 | 40 |
| 122.410 | | | 8.287 | | | 175.767 | 644 |
| 122.451 | | | 7.888 | | | 172.961 | 37 |
| 123.003 | | | 7.943 | | | 172.489 | 51 |
| 126.171 | | | 7.734 | | | 174.436 | 21 |

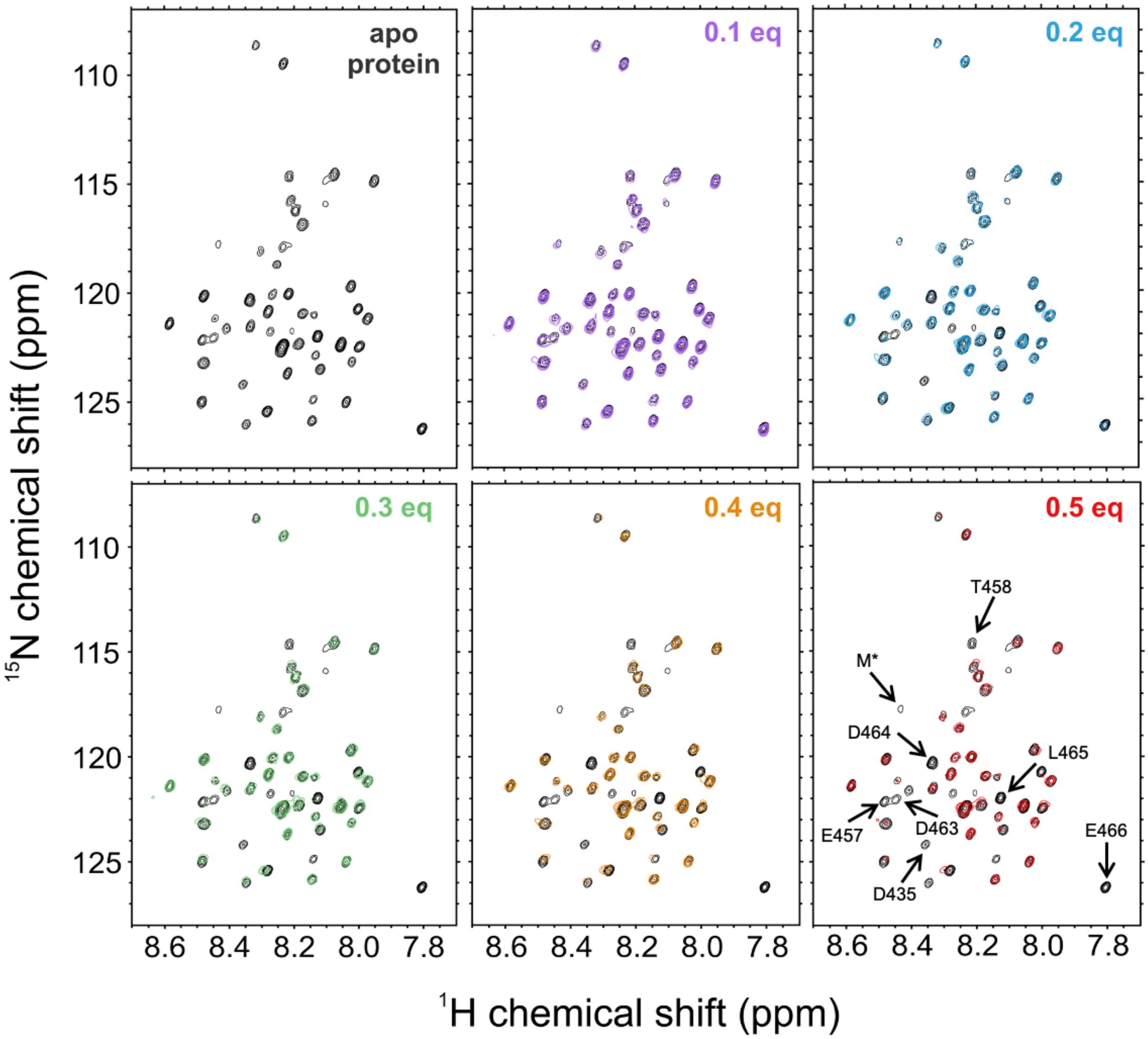


**Figure S10. NMR analysis of Cu(II) binding to VimT.** $^{1}$H–$^{15}$N HSQC spectrum of apo protein (38 μM) and spectral overlays of spectra recorded after the addition of 0.1, 0.2, 0.3, 0.4, 0.5 equivalents (eq) of Cu(II) at pH 6.5, recorded at 800 MHz $^{1}$H frequency. M* denotes the unremoved methionine residue from the purification 6xHis-tag that remains after TEV cleavage.

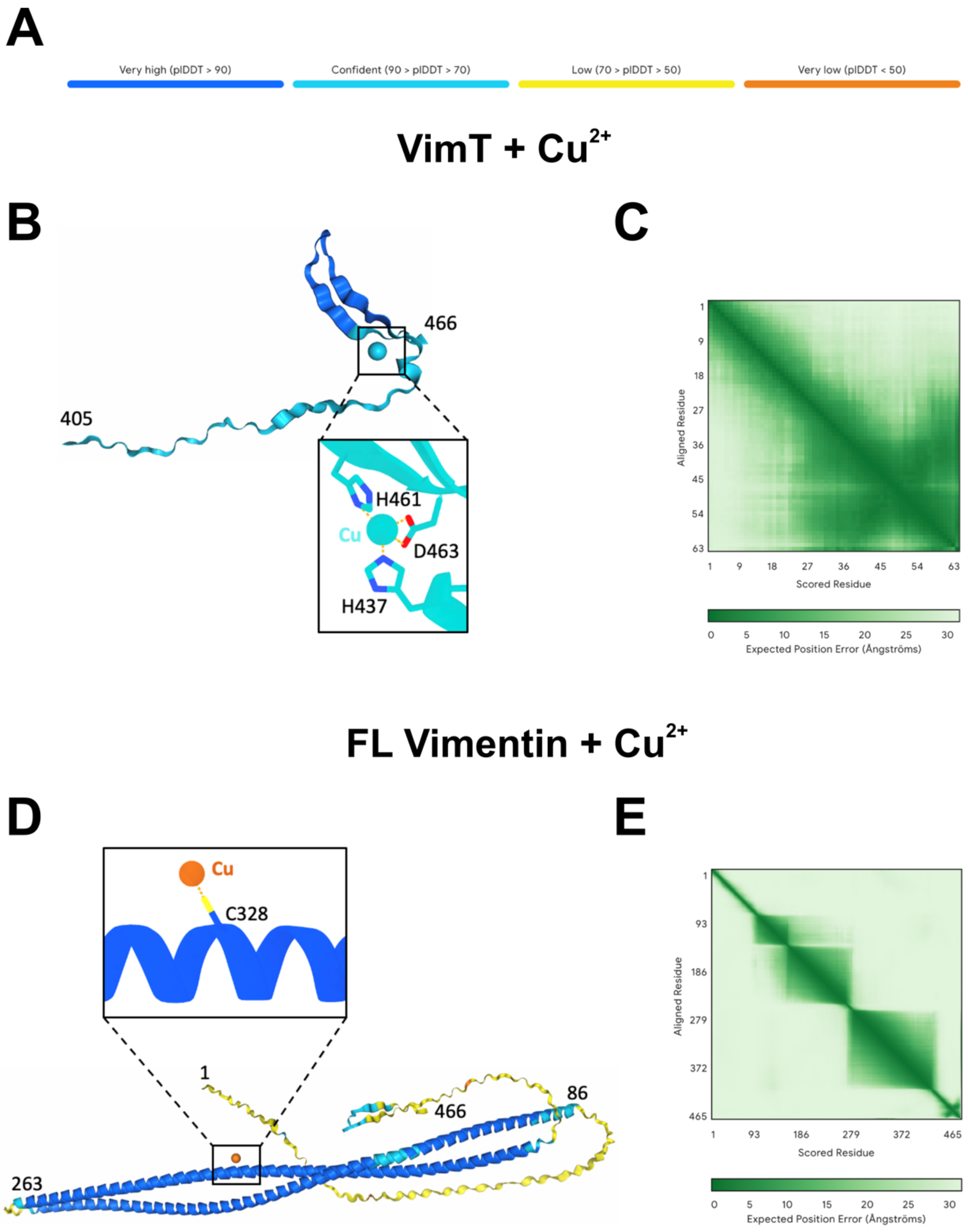


**Figure S11. AlphaFold 3.0[10] models for different forms of vimentin at varying ratios of protein to copper ($Cu^{2+}$) ions.** (A) The color-coded per-atom confidence scores (pLDDT). Note that as you go from the tail domain fragment (VimT) to the full-length (FL) and increasing copies of vimentin protein (not included), the pLDDT scores of the head and tail regions decrease to low (yellow) or very low (orange). The input of (B) VimT and $Cu^{2+}$ displays the copper ion coordinated to H461, D463, and H437, (D) FL vimentin (monomer) and $Cu^{2+}$ shows the copper ion coordinated to the single cysteine residue (C328) in the rod domain. (C, E) The corresponding predicted alignment error matrices for B and D, respectively. The darker green patches represent more confidence with larger distinct areas indicating more rigid units. The darker patches also highlight regions with predicted interdomain contacts.
